# Fractal triads efficiently sample ecological diversity and processes across spatial scales

**DOI:** 10.1101/2021.01.15.426868

**Authors:** Elizabeth G. Simpson, William D. Pearse

## Abstract

The relative influence of ecological assembly processes, such as environmental filtering, competition, and dispersal, vary across spatial scales. Changes in phylogenetic and taxonomic diversity across environments provide insight into these processes, however, it is challenging to assess the effect of spatial scale on these metrics. Here, we outline a nested sampling design that fractally spaces sampling locations to concentrate statistical power across spatial scales in a study area. We test this design in northeast Utah, at a study site with distinct vegetation types (including sagebrush steppe and mixed conifer forest), that vary across environmental gradients. We demonstrate the power of this design to detect changes in community phylogenetic diversity across environmental gradients and assess the spatial scale at which the sampling design captures the most variation in empirical data. We find clear evidence of broad-scale changes in multiple features of phylogenetic and taxonomic diversity across aspect. At finer scales, we find additional variation in phylo-diversity, highlighting the power of our fractal sampling design to efficiently detect patterns across multiple spatial scales. Thus, our fractal sampling design and analysis effectively identify important environmental gradients and spatial scales that drive community phylogenetic structure. We discuss the insights this gives us into the ecological assembly processes that differentiate plant communities found in northeast Utah.

## 1 Introduction

Ecological community assembly processes, such as environmental filtering (Kraft et al. 2015), competition (Mayfield and Levine 2010), dispersal (Vellend 2010), and facilitation (Valiente-Banuet and Verdú 2007), determine the diversity and structure of plant communities. Ecological processes operate at and across spatial scales: density-dependent biotic interactions tend to occur at local scales; environmental filtering often constrains species at community scales; and biogeographic processes define the species pool at regional to continental scales (Weiher et al. 1998; Cavender-Bares et al. 2009). These general trends simplify complex interactions between these processes that result in observable patterns, and many processes, such as dispersal, explicitly operate across multiple spatial scales (Chave 2013). Temporal scales similarly affects our understanding of process from pattern (Cavender-Bares et al. 2009; Wiens 2018) and at longer time-scales evolutionary processes — like selection, drift, and speciation — often play a key role in ecological assembly (Vellend 2010).

Phylogenetic diversity metrics represent the evolutionary history of an assemblage and provide good proxies of an assemblage’s ecological structure, despite known difficulties in inferring ecological processes from changes in phylogenetic pattern across environment (Webb et al. 2002; Cavender-Bares et al. 2009; Mayfield and Levine 2010; Mouquet et al. 2012). These metrics are affected by both the spatial grain, or sampled resolution, and spatial extent, or total study area of a study (Wiens 1989; Levin 1992; Rahbek 2005). Making the spatial grain and extend of a study larger increases (and eventually saturates) the number of species captured by that study (Crawley and Harral 2001; Adler et al. 2005; Fridley et al. 2005). For phylodiversity metrics, increasing spatial extent results in a larger species pool and phylogenetically clustered assemblages [where co-occurring species are more related than expected by chance, Cavender-Bares et al. (2006) and Swenson et al. (2006)]. Increasing a study’s spatial grain has a similar effect — assemblages shift from being overdispersed (containing species less related to one-another than expected by chance) to being clustered or phylogenetically random (Swenson et al. 2007). While these are general, and not universal, patterns, it is uncontroversial to state that a study’s spatial grain and extent affect observed diversity (Cavender-Bares et al. 2009; Vamosi et al. 2009; Pearse et al. 2013). However, it is often challenging to know the spatial scales that influence a system’s diversity patterns *a priori*, and thereby pick an appropriate grain and extent to best measure ecological processes of interest (Wheatley and Johnson 2009; Jackson and Fahrig 2015).

Fractal sampling designs provide a potential solution to the problem of knowing the appropriate spatial scale at which to measure biodiversity by systematically spacing sampling locations at in-creasingly closer or farther distances (Ewers et al. 2011; Marsh and Ewers 2013). Fractal sampling captures information more efficiently than grid or transect designs (Kallimanis et al. 2002) for a comparatively smaller time, effort, and financial input per sampling location than many other sampling strategies (Halley et al. 2004; Albert et al. 2010; Luzuriaga et al. 2012). Additionally, this design does not need to be oriented across a linear environmental gradient already known to affect diversity, making them useful for exploratory work in comparison with traditional straight-line transects (Marsh and Ewers 2013).

However, current fractal designs cannot be extended or intensified to include additional spatial scales while maintaining initial sampling locations. Given that we often do not know, *a priori*, the appropriate spatial scale for sampling, it would be valuable for a fractal sampling design to have the flexibility to add or exclude spatial scales as needed. We outline, in Figure 1, a equilateral-triangle-based fractal sampling design, whereby we nest additional fractals within an existing layout. This means that, Marsh and Ewers (*c.f.* 2013), we add two new points, not three, to nest triangles within each other. This allows us to intensify or expand the design as needed, to assess questions at different spatial scales, while maintaining temporal continuity among sampling locations.

**Figure 1:**
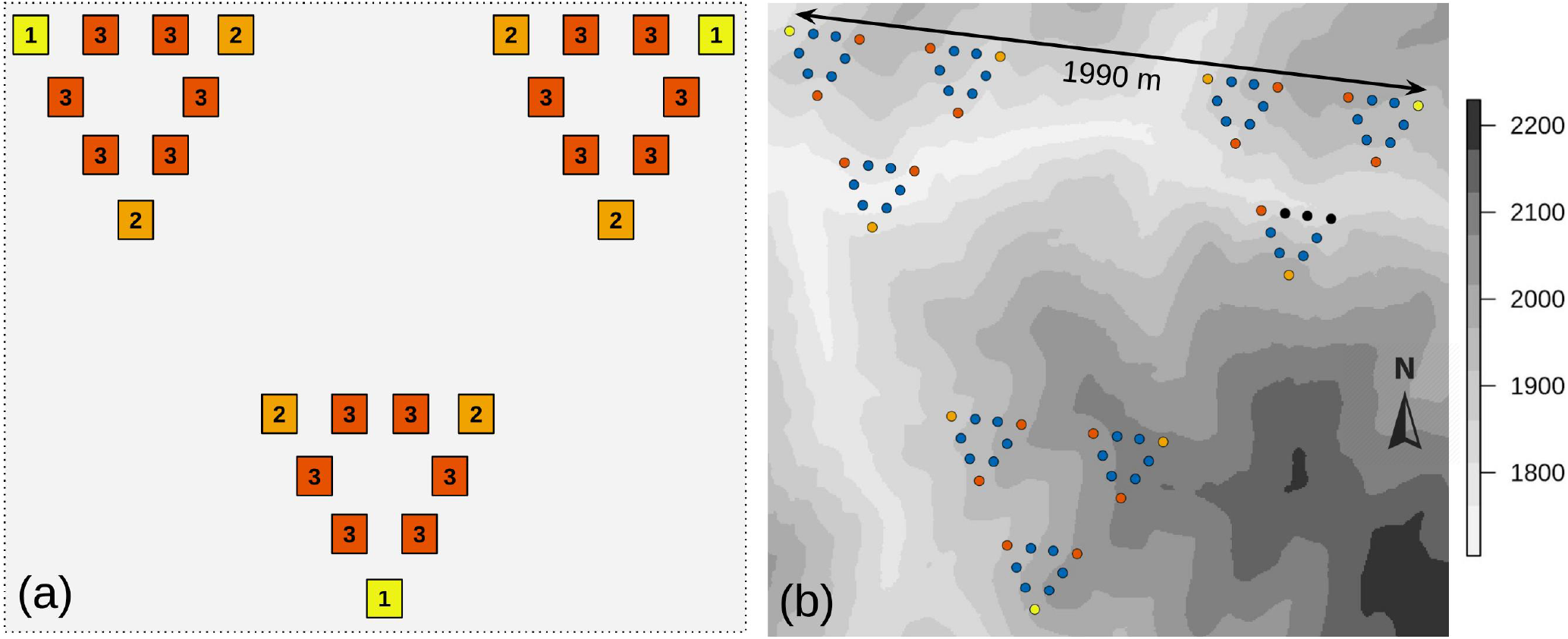
Overview of fractal sampling design. (a) A conceptual overview of how to build a fractal sampling design, in this case of three levels—‘triads’. First, choose three, initial plots ([1], yellow), at the vertices of an equilateral triangle that spans the greatest distance of interest at the study site. These sampling locations are the first and largest ‘triad’, and so the three plots in the first ‘triad level’. To build the second triad level, add two additional plots at the vertices of new equilateral triangles whose sides are 1/3 the length of the first triad. Critically, these new plots ([2], orange) are nested *within* the first triad level, and thus only two additional sites are needed because none of the outer site positions need be moved (*c.f.* Marsh and Ewers 2013). The third triad level ([3], dark orange) is analogously established within the second triad level. (b) Fractal sampling design applied at Right Hand Fork. We established and surveyed an initial set of three triad levels of plots in summer 2017 and re-surveyed them in 2018 (warm colors that match triad levels in (a). To assess whether we had sampled at a fine enough spatial scale to capture changes in diversity across environment, we added and surveyed a 4th triad level during summer 2018 (blue). The nested nature of our design allowed us to add these plots within the sampling arrangement, allowing us to continue monitoring from the third triad level sites. Distance between plots in the 1st, 2nd, 3rd, and 4th triad levels are 1990, 663, 221, and 74 meters respectively (*i.e.*, as in (a), each triad is nested in third). Due to safety concerns, we did not (re-)survey some plots in 2018 (black). Background grayscale shows elevation based on five-meter digital elevation model (Utah Automated Geographic Reference Center 2007).

We use this fractal sampling design to assess changes in plant phylogenetic and species diversity, and inferred ecological processes, across aspect and elevation in northeastern Utah. The multi-scale nature of this design allows us to couple this diversity-environment assessment with a variance components analysis to pinpoint the spatial scale(s) at which species and phylogenetic diversity varied most. We demonstrate the statistical power of fractally-nested designs across spatial scales, along with their ability to efficiently detect changes in diversity across environmental gradients and flexibility to address broader and finer spatial scales as needed.

## 2 Material and Methods

We aimed to test the ability of nested fractal sampling to quantify how phylogenetic and species diversity vary across environment, whether this variation is scale dependent, and at what scale that variation drives differences in diversity between assemblages. Below, we outline our approach to address each of these questions in turn. First, we demonstrated how a fractal sampling design provides more statistical power across spatial scales than random sampling. Then, we assessed how diversity metrics vary across elevation and aspect by surveying vascular assemblages using this sampling design in the field. Finally, we assessed whether spatial scale influences these metrics, by partitioning the variance associated with calculating that metric across the spatial scales in our fractal design. All software packages referenced below are for R (R Core Team 2020), and all data collected and code to reproduce analyses are openly released (Supplementary material Appendix 2, 3).

### 2.1 Study site, description, and survey methods

Our field site, located along the Right Hand Fork of the Logan River in Cache National Forest, UT (41.77003, - 111.59168), contains a variety of potentially interacting environmental gradients (Figure 1). The elevation spans 1719–2106 meters from riparian to ridge-line habitat. Numerous cliffs, rocks, and up to 54° slopes add fine-scale variation across the site. Overall vegetation type reflects aspect direction; sagebrush steppe on south-facing slopes and conifer forest on north-facing slopes (Lowry et al. 2007). Local land-use includes recreation along two trails that cross the site and permitted livestock grazing in about half of the plots (USDA Forest Service 2018).

We determined sampling location coordinates *a priori* at our site using a fractal sampling design (Figure 1) and navigated to these locations using a GPS, accurate to within 10 meters. At each 1 m^2^ plot we comprehensively surveyed each vascular plant species’ percent canopy cover by dividing each plot into four quadrants and using a 10 x 10 0.25 m^2^ grid to standardize cover estimates. Plants were identified using local herbarium resources, identification experts, and field guides. During June-August 2017, we established and surveyed 27 plots in three triad levels at 1990, 663, and 221 meters apart. During June-October 2018, we added an additional 54 plots in a 4th triad level at 74 meters apart) and surveyed all 78 plots. Due to safety concerns (the sites were on or close to cliffs), we did not survey 3 of the 81 plots in 2018. We report here results from the 2018 survey, but release the surveys, sampling locations, and meta-data for both the 2017 and 2018 surveys along with replicated analysis for the 2017 data (Supplementary material Appendix 2,3). All trends are qualitatively identical between the two surveys (Figures 3 and Supplementary material Appendix 1 Figure A2). To represent each plot’s topography, we measured aspect (in degrees, converted to a north-south gradient using a cosine function) using a compass, the slope (in degrees, average of uphill and downhill from the plot) using a clinometer, and the elevation (in meters) using the altimeter in a GPS.

### 2.2 Overview of our nested fractal sampling design

We outline our sampling design here, and in Figure 1. First, we placed three sampling locations at the vertices of an equilateral triangle whose side length spanned the spatial extent of the study area (c. 1990 meters). From each of the points, we added two additional sampling locations at the vertices of three new equilateral triangles whose sides were 1/3 the length of the first triad. We continued to nest sampling locations inward to add a third and (in 2018) a fourth triad level. By only adding two sites as each triad level (spatial scale) is added, instead of three (*c.f.* Marsh and Ewers 2013, where each successive triad is centered at what would be the higher level’s site), we saved 3^1^ plots for the 2nd triad level, 3^2^ plots for the 3rd triad level, and thus when we added a fourth triad level (in 2018) to our existing field system we saved 3^3^ (27) plots. The improved the efficiency of our fractal sampling design gave us temporal continuity in sampling locations as we investigated a finer spatial scale in our study area.

Here we provide a brief overview of how our sampling design concentrates plot comparisons across spatial scales to effectively address multi-scale questions; see Marsh and Ewers (2013) for a formal review of the statistical power of fractal designs. In Figure 2, we compare the pairwise distances among plots for fractal designs (in red and blue) with a distribution of randomly-placed designs (in grey). Fractal designs concentrate pairwise distances (or comparisons) of plots at specific spatial scales (in red, Figure 2), sacrificing comparability (and so statistical power) at some distances (in blue, Figure 2). This maximizes information content across all the spatial scales within the study’s spatial extent. Conversely, random sampling designs diffusely compare sites across spatial scales, concentrating information at the median spatial distance within the study’s spatial extent.

**Figure 2:**
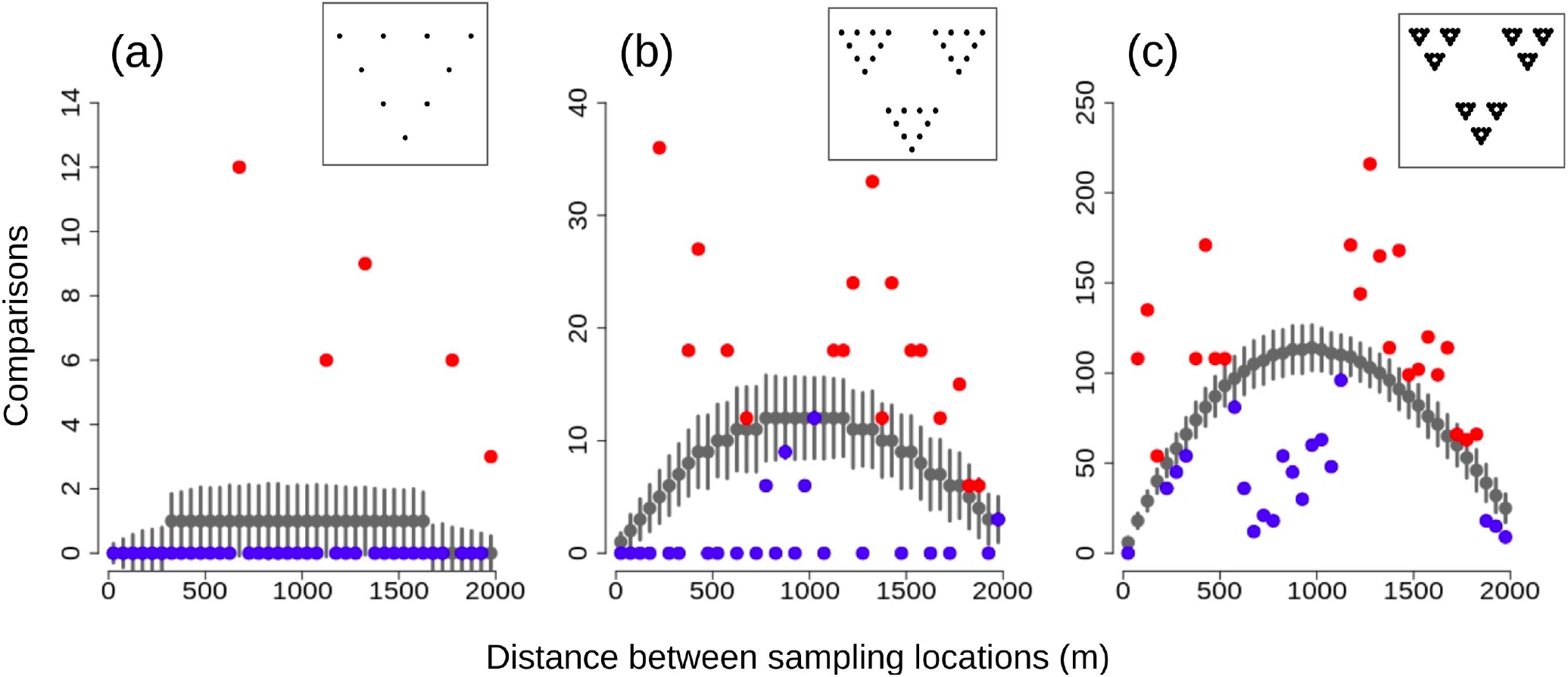
Our fractal sampling design concentrates statistical power across spatial scales (red dots) compared to random sampling designs with the same number of sampling locations (gray dots). (a) 2-triad, 9 plot, (b) 3-triad, 27 plot, and (c) 4-triad, 81 plot fractal sampling designs, built as described in Figure 1, all show the number of comparisons possible across a maximum sampling distance of 1900 meters for a fractal sampling design (red and blue dots) compared to the distribution of comparisons possible for 1000 randomly chosen sampling locations (gray dots, 95% confidence intervals shown as bars). Points in red demonstrate where there are more comparisons for a given distance class than the random sampling design distribution; (a) 5/9 plots, (b) 13/27 plots, and (c) 20/81 plots. Conversely, points in blue demonstrate fewer comparisons for a given distance class than the distribution of random sampling locations. The layout of the fractal sampling points is embedded in the upper right corner of each plot.

### 2.3 Diversity across environment

We calculated the diversity of the plant assemblage at each plot using two species-level and three phylogenetic diversity metrics that provide different insights about community structure and the potential drivers of community assembly (Tucker et al. 2017). Our first two metrics, species richness and Faith’s PD (Faith 1992), both capture the richness of diversity at each site. Faith’s PD correlates with species richness because it adds the phylogenetic branch lengths of all species present in a community, however it often gives more information about a community because it accounts for relatedness among species. Other metrics—*SES*_*MPD*_ and *SES*_*MNTD*_—place the relative phylogenetic divergence of species at a site in the context of the wider species pool (Webb 2000; Kembel 2009). This provides a specific context for how the each sampled community may have been assembled from this possible species pool, as opposed to drawing from a larger phylogeny which may include taxa that are not relevant to the sampled community. We also assessed a standard metric of diversity, Simpson’s diversity index (Simpson 1949), to test the effectiveness of this metric to represent changes in diversity across environment. All metrics were calculated using pez (Pearse et al. 2015), picante (Kembel et al. 2010), vegan (Oksanen et al. 2019), and the phylogenetic tree generated from Zanne et al. (2014) using pez::congeneric.merge. We modeled community variation across gradients using an additive linear model of each diversity metric across aspect and elevation for all 78 surveyed plots. To test the ability of our design to detect changes in diversity-environment relationships at different spatial scales, we re-fit these models using only the 26 plots from the 3rd triad level and only the 8 plots from the 2nd triad level.

### 2.4 The effect of spatial scale on diversity

We assessed whether our design captured different information at different spatial scales, using a variance components analysis to contrast how variance partitions across our nested triads. We calculated the amount of variation in each diversity metric attributable to a given triad level in our fractal design using variance components analysis following Crawley (2012). We fit a Bayesian linear hierarchical model with default priors using rstanarm (Goodrich et al. 2020), structured to sequentially partition the variance present in the modeled diversity metric from the largest (first) triad through to the smallest (fourth) triad. We fit our model in a Bayesian rather than frequentist framework to avoid singular fits associated with fitting the largest triad level, which contains only three groups. To ensure that our Bayesian approach to estimating variance was robust, we compared our observed data to underlying data whose nested structure was randomly broken. Thus, in 999 bootstrap randomizations, we randomly permuted sites’ locations and performed the same variance components analysis. We ranked our observed (real, unpermutted) data within these bootstrap randomization, significant at *α* = 0.05, to statistically test whether each biodiversity metric showed an unexpected spatial pattern at that triad level.

## 3 Results

We used our fractal sampling design to assess changes in biodiversity metrics across environment and differing spatial scales. We identified a total of 120 species within our plots at RHF and surveyed a mean of 11 species/plot within a range of 5 to 21 species/plot. All phylogenetic diversity metrics (PD, *SES*_*MPD*_, and *SES*_*MNTD*_) varied significantly across aspect; *SES*_*MPD*_ also varied across elevation (Figure 3). For *SES*_*MNTD*_, sampling at the 2nd and 3rd triad levels (*i.e.*, with only 8 and 26 sites) would have been sufficient to detect these relationships in the 2018 survey (Figure 3). We only needed the 3rd triad level to detect changes in *SES*_*MPD*_ and Faith’s PD for the 2018 survey as well. We found similar trends in the 2017 data, detecting changes in *SES*_*MNTD*_ and Faith’s Pd across aspect at the 3rd triad level (Supplementary Material Appendix 1 Figure A2).

**Figure 3:**
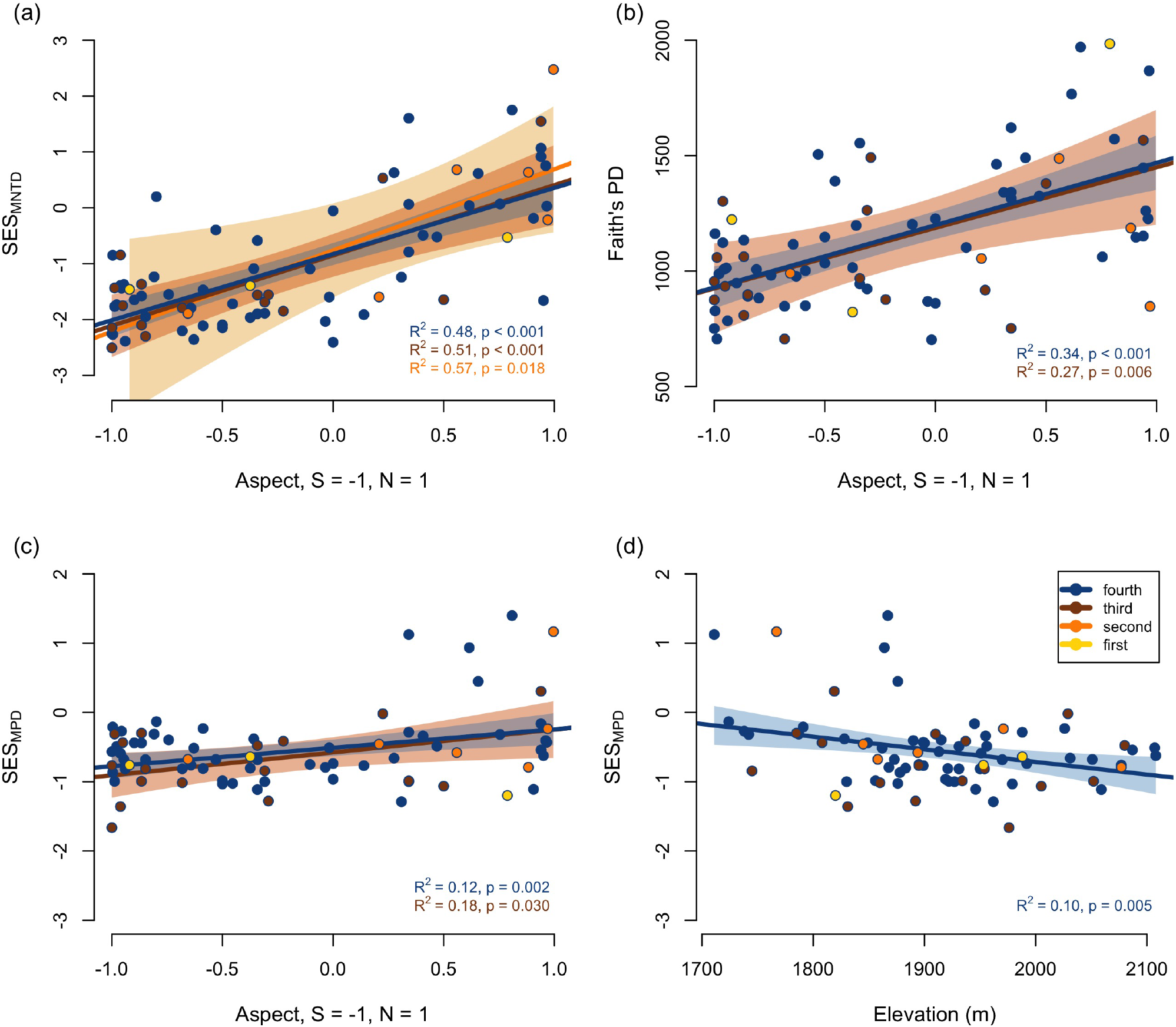
Changes in phylogenetic diversity across environment detected at different spatial scales by our fractal sampling design. (a) *SES*_*MNTD*_, (b) Faith’s PD, and (c) *SES*_*MPD*_ were greater on more northern aspects, and (d) *SES*_*MPD*_ decreased as elevation increased. While models of diversity across environment were tested for all triad levels (see Fig. 1) only significant models are plotted (with 95% confidence intervals). We detected a change in *SES*_*MNTD*_ aspect and the 4th–2nd triad level and a change in *SES*_*MPD*_ across aspect at the 4th and 3rd triad level. While changes in Faith’s PD across aspect and *SES*_*MPD*_ across elevation were detectable only at the finest sampling of the fourth triad level *SES*_*MNTD*_ was more sensitive and thus able to detect changes with less sampling. Points are color-coded based on their triad level; blue is the 4th level with 78 surveyed locations, dark orange is the 3rd level with 26 surveyed locations, light orange is the 2nd level with 8 surveyed locations, and yellow is the 1st level with 3 surveyed locations.

For each diversity metric calculated, we used a variance components analysis to assess the variance associated with each spatial scale in our fractal sampling design (Figure 4). Species richness and Faith’s PD significantly associated with the largest, 1st triad level, accounting for 75% and 84% of the variance in each of these metrics. Additionally, Faith’s PD significantly associated with variance in both other triads, 2% and 11% of the variance at the 2nd and 3rd levels respectively. In a similar pattern, species richness associated with 6% of the variance at the 3rd triad level. Both *SES*_*MNTD*_ and *SES*_*MPD*_ pick up larger amounts of variance across spatial scales.They account for 27% and 34% of variance (*SES*_*MNTD*_) and 16% and 16% of variance (*SES*_*MPD*_) at the variation 2nd and 3rd triad levels respectively.

**Figure 4:**
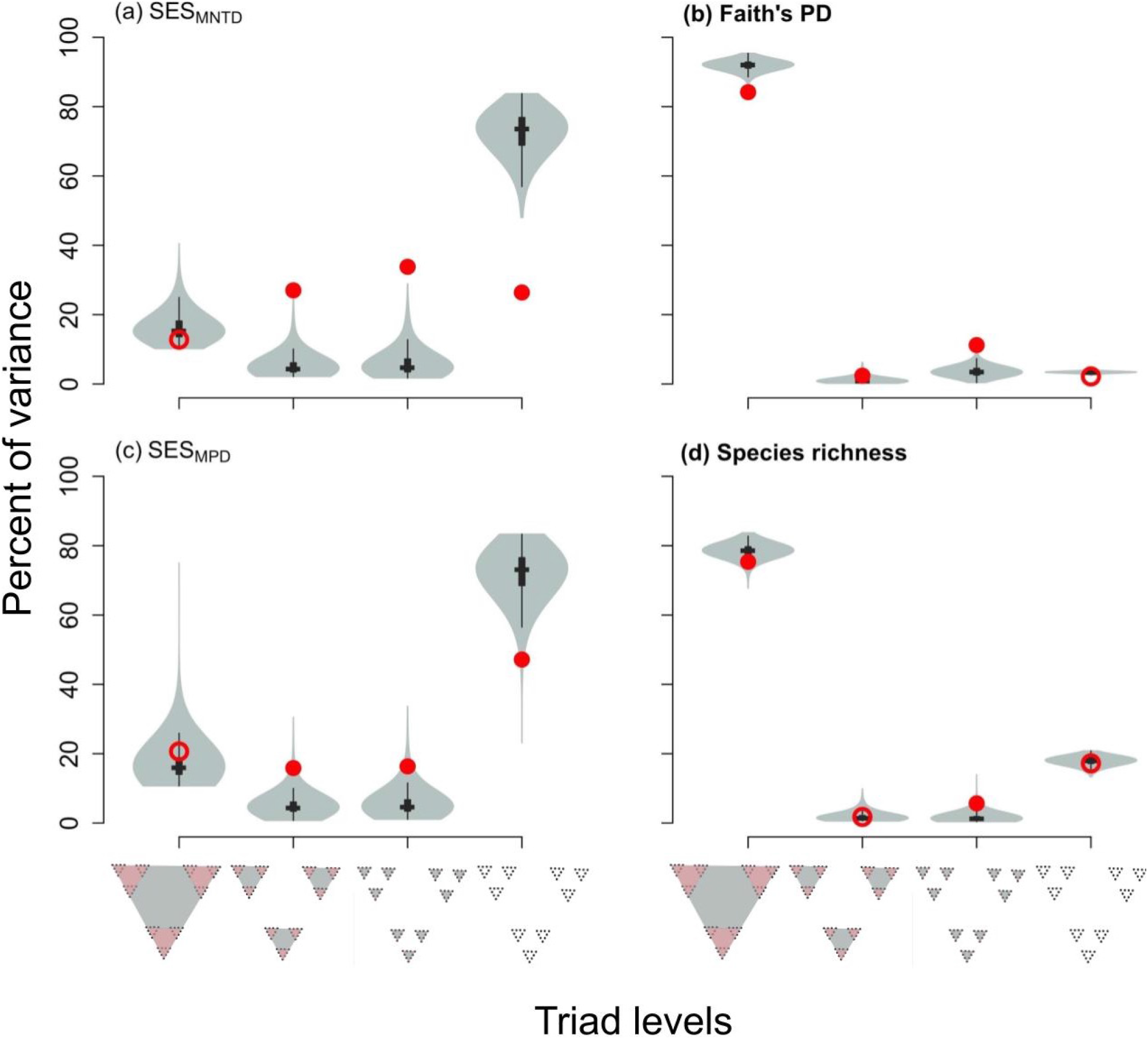
Spatial scales, represented by the fractal sampling design, capture variance in phylodiversity metrics. For each diversity metric, (a) *SES*_*MNTD*_, (b) Faith’s PD, (c) *SES*_*MPD*_, (d) species richness, the red dots show the percent of variance captured by each successive spatial scale in our fractal sampling design, calculated using a variance components analysis. Randomized diversity values for each triad level (999 iterations) shown as gray violin plots. Filled red circles indicate percent of variance values that are significantly different from the randomized diversity variance at each triad level (violin plots). These spatial scales directly account for a portion of the variance in each of these diversity metrics. P-values for calculated percentages of variance that are significantly different from the random distribution of potential variation captured by each triad level: *SES*_*MNTD*_ (2*nd* = 0.002, 3*rd* < 0.001, 4*th* = 0.001, Faith’s PD (1*st* = 0.001, 2*nd* = 0.048, 3*rd* < 0.001), *SES*_*MPD*_ (2*nd* = 0.010, 3*rd* = 0.021, 4*th* = 0.004), species richness (1*st* = 0.026, 3*rd* = 0.011).

## 4 Discussion

Our fractal sampling design captured empirical changes in multiple plant biodiversity metrics across different environmental gradients and spatial scales. Among the phylogenetic diversity metrics we calculated, Faith’s PD, a metric that includes information about the overall evolutionary history of species in a community, detected the most information about community differences, at the largest spatial scale we studied. Conversely, *SES*_*MNTD*_, a metric that focuses on more recent evolutionary history, detected the most information about how assemblages change at smaller spatial scales. Below, we discuss how this design, coupled with modeling change across environment and a variance components analysis, provides a practical and effective way to assess how diversity and inferred ecological processes change across space and environment.

### 4.1 Abiotic conditions dominate broad-scale assembly

We predominantly detected changes in assemblage structure across aspect. We show shifts from phylogenetically clustered assemblages (containing closely related species) on south-facing slopes (less PD, negative *SES*_*MNTD*_ and *SES*_*MPD*_; Figure 3) to more distantly related assemblages on north-facing slopes (more PD, near-zero to positive *SES*_*MNTD*_ and *SES*_*MPD*_; Figure 3). Studies of species diversity across aspect find that communities on south-facing slopes tend to contain fewer species than north-facing slopes (Cantlon 1953; Olivero and Hix 1998; Fridley 2009), while we found no difference in the number of species on opposing slopes. However, they also find that south-facing assemblages tend to have more consistently similar species compositions, compared to north-facing assemblages.

In the Northern hemisphere, greater sun exposure on south-facing slopes intensifies heat, plant tissue damage, and reduces soil moisture in an already arid climate (Lowry et al. 2007), which likely limits the number and type of species able to grow and persist (Keddy 1992; Weiher et al. 1998). Our phylogenetic diversity metrics align with this constraint leading to lower phylogenetic diversity (less PD) on south-facing slopes. Additionally, that the metrics we calculated that account for species richness (*SES*_*MNTD*_ and *SES*_*MPD*_) change across this gradient demonstrates that environment constrains phylogenetic diversity to clades whose members can tolerate these conditions.

Conversely, north-facing slopes receive less sun exposure which results in cooler temperatures and better soil moisture retention — a more favorable set of growth conditions in an otherwise resource-limited environment (Moeslund et al. 2013). The phylogenetic clustering we observe on south-facing slopes does not inherently indicate environmental filtering (Mayfield and Levine 2010). However, we observe changes in diversity across environment aligning with studies that demonstrate that environment constrains (and thereby filters) phylogenetic (Webb 2000; Helmus et al. 2007), species (Luzuriaga et al. 2012; Laliberté et al. 2014), and functional diversity (Luzuriaga et al. 2012; Maire et al. 2012; Bello et al. 2013).

Change in phylogenetic structure across environment at Right Hand Fork provides evidence that environmental filtering plays a role in community assembly across these plots. We recognize that we have not experimentally quantified whether species presence or absence relies solely on abiotic conditions (as is necessary to prove environmental filtering; Kraft et al. 2015), but we do show that changes in environment map onto changes in ecological communities. By combining our spatially explicit structure of our sampling design with a variance components analysis, we can, however, precisely pinpoint the spatial scales at which environment is likely to be structuring community assembly. For Faith’s PD and species richness, the largest spatial scale (1st, spaced at 1990 meters), captured the most variance in these metrics (84% and 75% respectively). Surprisingly, however, these values are significantly less variation than our null expectations, and we suggest this surprising result stems from two opposing forces. First, species richness (and so Faith’s PD, which is often correlated with it, Tucker et al. (2017).) is likely driven by processes such as lineage diversification that operate across broader spatial scales than we measure here. We are currently extending the sampling of our fractal system further in an attempt to capture additional processes operating across ecological timescales. Second, while these metrics are less sensitive to finer-scale processes than our other metrics (see below), they do still detect some pattern, thus reducing the variance explained at the broadest scale.

The similarity in variance partitioning patterns between Faith’s PD and species richness shows that generally speaking, they represent similar information about communities in this system (Tucker and Cadotte 2013). However, we were able to detect changes across environment with Faith’s PD but not species richness. This, coupled with our fractal design’s ability to capture slightly more variance in Faith’s PD than species richness (9%), supports the use of phylogenetic diversity as a more predictable and informative metric about assemblage composition.

### 4.2 Small-scale biotic assembly

Within the context of broad-scale assemblage differences driven by aspect, we found evidence for differences in biotic interactions at more local scales that demonstrate further community differentiation. Phylogenetic diversity metrics that account for the source pool of potential species (and *SES*_*MPD*_), capture variance across multiple, more local scales. Both *SES*_*MPD*_ and *SES*_*MNTD*_ detected the most variation in assemblage structure at finer scales (*SES*_*MND*_, 2nd and 3rd triad level, 27% and 34% respectively, *SES*_*MPD*_, 2nd and 3rd triad level, 16% and 16% respectively; Figure 4)). Since these phylodiversity metrics are calculated using a source pool of potential species, they account for broad-scale structure when assessing local context (Webb et al. 2002; Kembel 2009), unlike our other metrics. We suggest this makes these metrics more sensitive to differentiation at and across local spatial scales, giving us a more nuanced picture of local variation in diversity. Perhaps most striking, *SES*_*MNTD*_ demonstrates strong spatial structure at the middle two scales (2nd and 3rd triad) in our sampling design, accounting for close to 2/3 of the variance in this metric. The Brownian motion model of trait evolution assumed by many studies of phylogenetic assemblage more strongly predicts that close-relatives’ traits (Letten and Cornwell 2015). This pattern of *SES*_*MNTD*_ being more strongly predictable than *SES*_*MPD*_ likely stems from the inherently greater predictability of close-relatives’ niches under such models. This insight, along with assumed phylogenetic conservatism, supports *SES*_*MNTD*_ as a strong diversity metric to detect assemblage differences at and across the local spatial scales we assessed at Right Hand Fork.

### 4.3 Conclusion

We conclude that changes in phylogenetic diversity and inferred ecological process across environment and spatial scale can be efficiently detected using a fractal design and variance components analysis. Phylogenetic diversity metrics gave us more information about assemblage composition than species richness alone. Faith’s PD accounted for broader patterns of species presence in response to overall environment, while and *SES*_*MPD*_ reflected how biotic interactions generate localized environmental heterogeneity. Our spatially explicit design allows systematic comparison of patterns and hypotheses at multiple spatial scales. An advantage of our fractal approach is that it is impartial with regard to any particular environmental gradient, and can be intensified and extended after establishment, which we leveraged to examine variation at a smaller spatial scale than initially sampled. This flexibility allows us to continue to investigate questions about the relationship between diversity and environment and the way spatial scale affects those relationships. For example, this sampling framework could be extended to study other drivers of community assemblage across a landscape such as soil temperature and texture. Systematic exploration of this system via a fractal sampling design will continue to allow us to investigate diversity across scale and environment using this powerful and efficient sampling design.

## Declarations

## Acknowledgements

We would like to thank the members of the Pearse Lab for their feedback and support and Mary Barkworth and Michael Piep for extensive assistance with learning to identify Utah plants.

## Statement of Authorship

All authors contributed to data collection, analysis, and manuscript preparation but EGS did the majority of data collection.

## Data accessibility

All data released in supplementary materials.

