## Appendix 1 for "Fractal triads efficiently sample ecological diversity and processes across spatial scales"

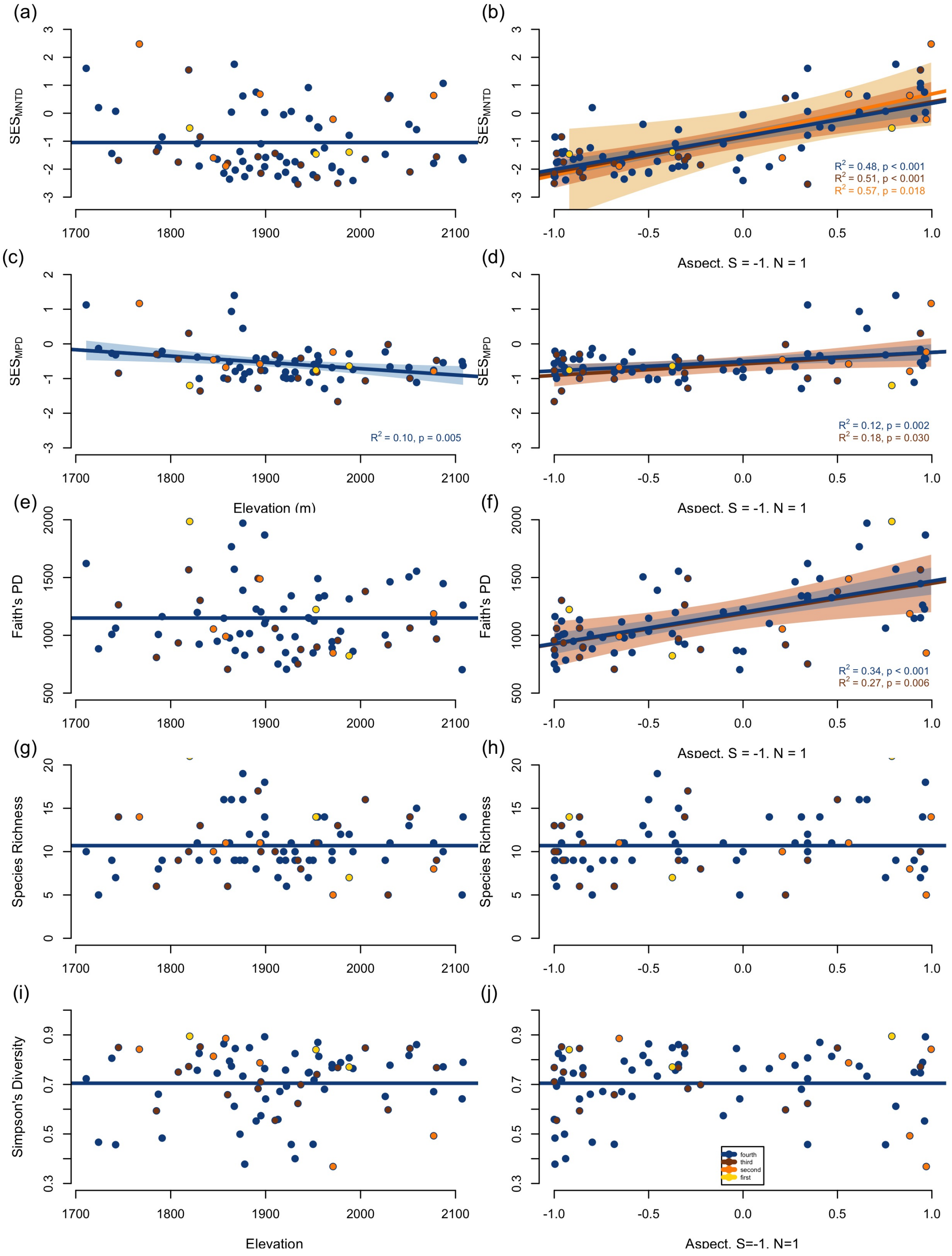


Figure A1: **All changes in diversity across environments detected at different spatial scales by our fractal sampling design.** Left column shows change (or lack thereof) in diversity across elevation while the right column shows the same diversity metrics across aspect (a, b) $SES_{MNTD}$, (c, d) $SES_{MPD}$, (e, f) Faith's PD, (g, h) species richness, and (i, j) Simpson's diversity. Color scheme and models plotted as described in Figure 3 caption.

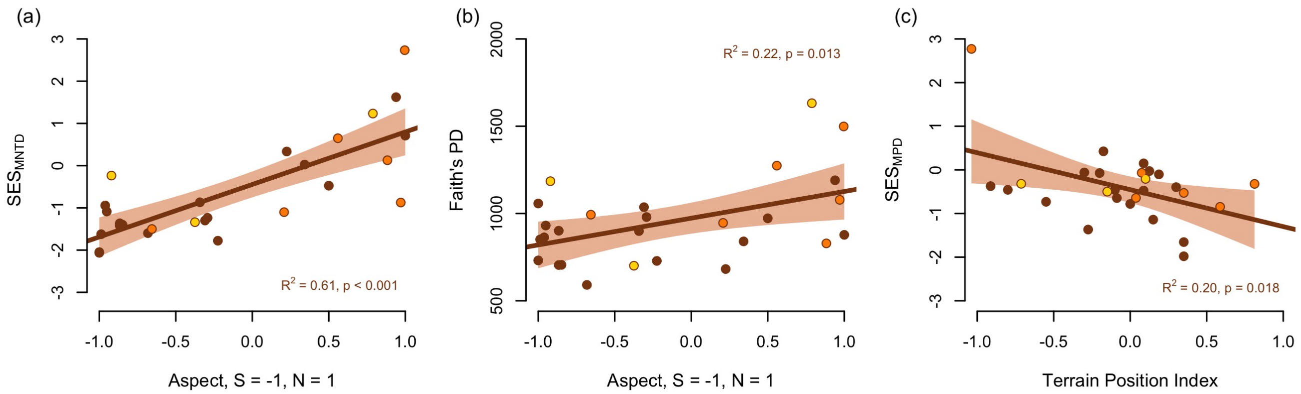


Figure A2: **Significant changes in diversity across environments detected at different spatial scales by our fractal sampling design, surveyed in 2017.** Each panel shows change in (a) $SES_{MNTD}$ across aspect, (b) Faith's PD across aspect, and (c) $SES_{MPD}$ across Terrain Position Index. Color scheme and models plotted as described in Figure 3 caption.

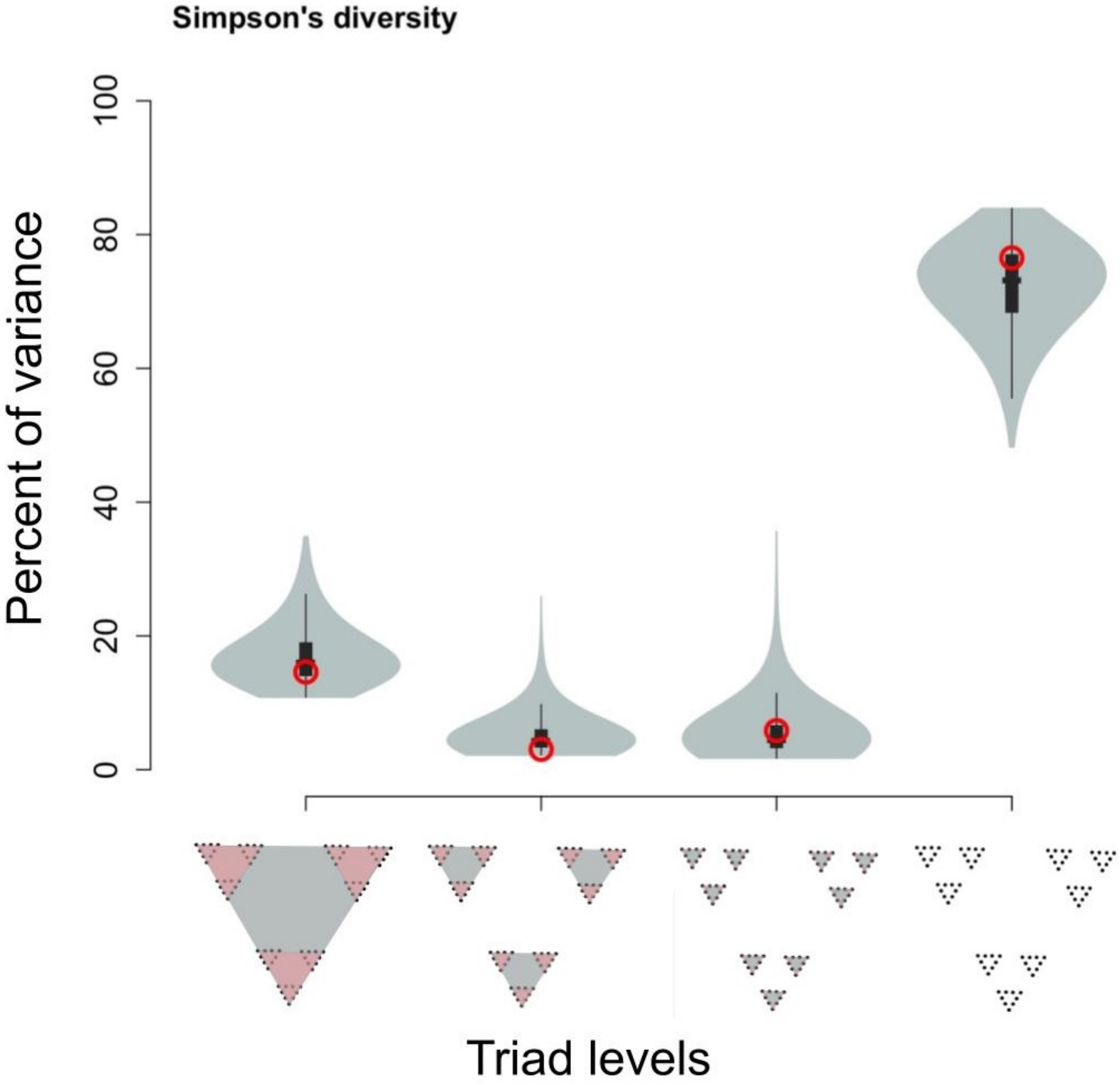


Figure A3: **Spatial scales, represented by the fractal sampling design, do not capture significant variance in Simpson's diversity.** Explanation of plot in text (Figure 4 caption.)

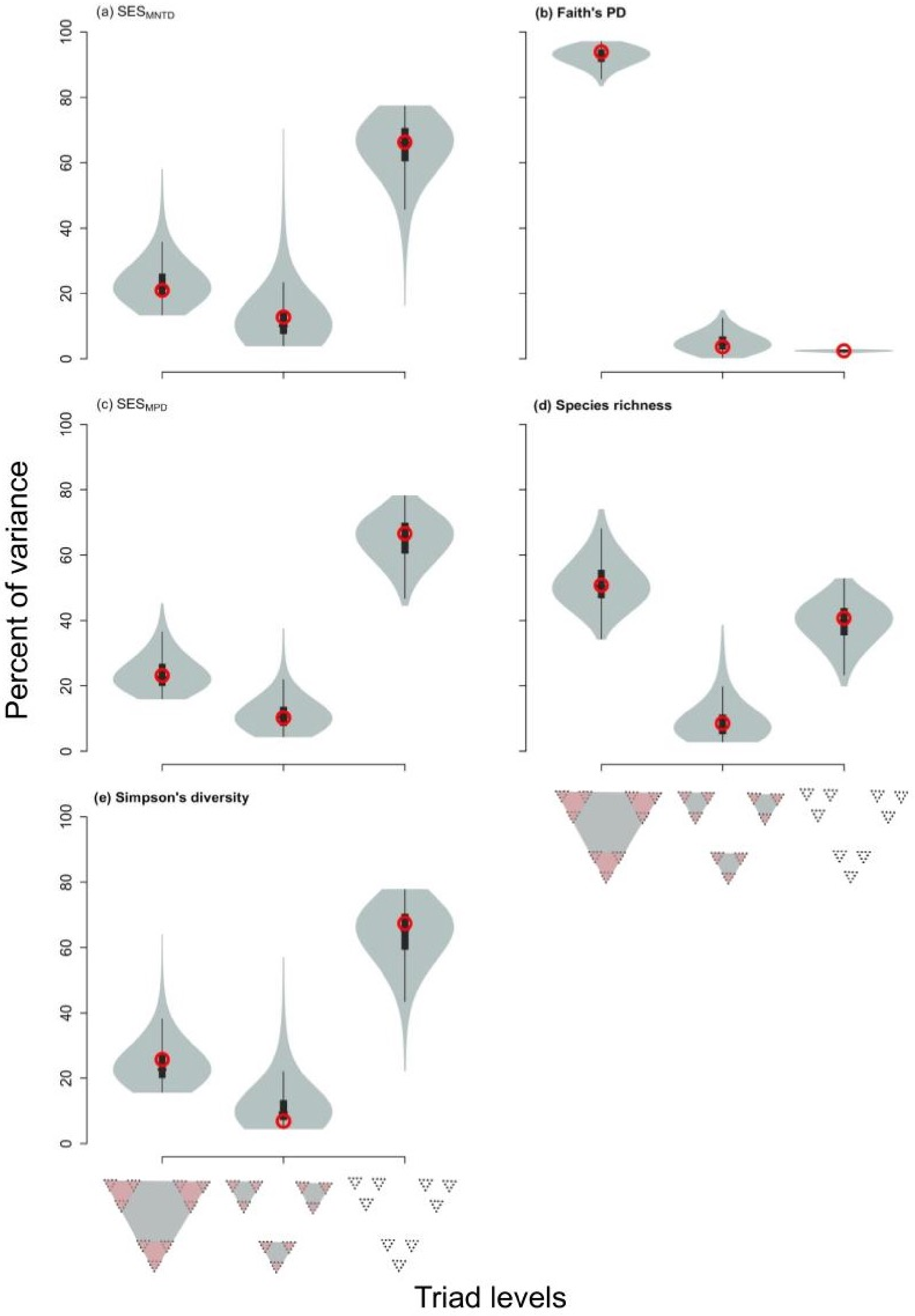


Figure A4: **Only three spatial scales, represented by the fractal sampling design, do not capture significant variance in any of the diversity metrics calculated from the 2017 survey data.** Explanation of plot in text (Figure 4 caption.)
