## Appendix 2 for "Fractal triads efficiently sample ecological diversity and processes across spatial scales": comm-diversity-across-env-and-scale-2017-supplement.R

```
# Assessing community (phylogenetic) diversity across environment - 2017 data
# Elizabeth Simpson & Will Pearse - 2017-9-21
#setwd("~/Documents/projects/fractal_sampling_div_rhf/analysis_code") # uncomment to set workind directory on computer

###########
# HEADERS #
###########
library(pez) 
library(vegan)
library(lme4)
library(visreg) # need to load this package to plot loaded .Rdata

# functions for splitting the data into different triad levels
subset.first.level <- function(data){
  one <- matrix(nrow=0,ncol=ncol(data))
  colnames(one) <- colnames(data)
  for (i in seq_len(nrow(data))){
    if (data$first[i] == data$second[i] & data$second[i] == data$third[i]){
      one <- rbind(one, data[i,])
    }
  }
  return(one)
}

subset.second.level <- function(data){
  two <- matrix(nrow=0,ncol=ncol(data))
  colnames(two) <- colnames(data)
  for (i in seq_len(nrow(data))){
    if (data$second[i] == data$third[i]){
      two <- rbind(two, data[i,])
    }
  }
  return(two)
}  

subset.env.17 <- function(data){
  three <- matrix(nrow=0,ncol=ncol(data))
  colnames(three) <- colnames(data)
  for (i in seq_len(nrow(data))){
    if (data$third[i] == data$fourth[i]){
      three <- rbind(three, data[i,])
    }
  }
  return(three)
}  

r2_and_p <- function(model){
  return(c(round(summary(model)$r.squared,2), round(pf(summary(model)$fstatistic[1],summary(model)$fstatistic[2],summary(model)$fstatistic[3],lower.tail=FALSE),3)))
}
##############################################################################
# Make comparative community object from raw precense & percent cover data ###
##############################################################################

################
# Community data 

# load in cover data, reformat, remove NAs, remove species essentially there as notes
data <- read.csv("../data/rhf_2017_cover.csv", as.is=TRUE)
comm <- with(data, tapply(Cover, list(Site,Species), function(x) mean(x,na.rm=TRUE)))
comm[is.na(comm)] <- 0
comm <- comm[,!grepl("^[a-z]+", colnames(comm))]
comm <- comm[,!grepl("\\(|/", colnames(comm))]
comm <- comm[,!grepl("sp\\.", colnames(comm))]
colnames(comm) <- tolower(colnames(comm))

################################
# Phylogeny (Zanne et. al. 2014) 

# load tree, reformat sp names, create combined data object
tree <- read.tree("../data/Vascular_Plants_rooted.dated.tre")
tree$tip.label <- tolower(gsub("_", " ", tree$tip.label))
tree <- congeneric.merge(tree, colnames(comm), split = " ")

####################
# Environmental data

# load in terrain variables, name rows with plot ID's, create comparative community object
env <- read.csv("../data/clean_rhf_terrain_for_2017.csv", as.is=TRUE)
env <- env[,3:10]
env <- subset.env.17(env)
rownames(env) <- with(env, paste0(first, second, third))

c.data <- comparative.comm(tree, comm, env=env)

###################################
### Calculate diversity metrics ###
###################################

# Calculte divresiy metrics from abundance community data - 
ses.mpd <- .ses.mpd(c.data)
ses.mntd <- .ses.mntd(c.data)
pd <- .pd(c.data)
simps.div <- diversity(c.data$comm, index="simpson")

# Compile metrics into a data frame with environmental information
data <- cbind(c.data$env, ses.mpd$ntaxa, ses.mpd$mpd.obs.z, ses.mntd$mntd.obs.z, pd[,1], simps.div)
data <- data[,c(1:3,5:13)]
colnames(data) <- c("first", "second", "third", "elev", "aspect", "slope", "tpi", "ntaxa", "sesmpd", "sesmntd", "pd", "simpsdiv")

##################################################################
# Load to here for VCA ###########################################
##################################################################

# Subset the data into different triad levels
one <- subset.first.level(data)
two <- subset.second.level(data)
# three is the whole data frame of all 26 surveyed plots

#########################################
# Modeling diversity across environment #

# Species richness
nsp3.all <- with(data, lm(ntaxa ~ elev + slope + aspect + tpi)) # no sig.
nsp <- with(data, lm(ntaxa ~ 1))

# SES_mpd (z) 
mpd3.all <- with(data, lm(sesmpd ~ elev + slope + aspect + tpi))
mpd3.tpi <- with(data, lm(sesmpd ~ tpi))

# SES_mntd (z) 
mntd3.all <- with(data, lm(sesmntd ~ elev + slope + aspect + tpi)) 
mntd3.a <- with(data, lm(sesmntd ~ aspect))

# Faith's PD
pd3.all <- with(data, lm(pd ~ elev + slope + aspect + tpi)) # aspect sig.
pd3.a <- with(data, lm(pd ~ aspect))

# Simpson's Div ####
simps3.all <- with(data, lm(simpsdiv ~ elev + slope + aspect + tpi)) # no sig
simps <- with(data, lm(simpsdiv ~ 1))

# output
output <- rbind(r2_and_p(mpd3.tpi), r2_and_p(mntd3.a), r2_and_p(pd3.a))
rownames(output) <- c("mpd3.tpi", "mntd3.a", "pd3.a")
colnames(output) <- c("r_squared", "p_value")

write.csv(output, "./figures-results/comm-2017-results.csv")

##################################################################################################################################################
# Plot Figure 3 ##############################################################################################################################
#setwd("~/Documents/projects/fractal_sampling_div_rhf/analysis_code/figures")
jpeg("./figures-results/fbsd-fig3-2017-supplement.jpeg", width = 9, height = 3, units = 'in', res = 300) # Make plot
par(mfrow=c(1,3))
par(mar=c(4,4.5,3,1))
par(oma=c(.7,.7,.7,.7))

# SESmntd ~ aspect
mntdCIa <- visreg(mntd3.a, plot=FALSE)
plot(mntdCIa, overlay=TRUE, line=list(lwd=0),
     fill=list(col=c("#D55E006E")),
     points=list(cex=0), xlab= "Aspect, S = -1, N = 1", ylab=expression(SES[MNTD]), legend=FALSE, cex.lab=1.1, ylim=c(-3.2, 3), axes=FALSE)
abline(mntd3.a, col= "chocolate4", lwd=3)

with(data, points(sesmntd ~ aspect, pch=19, cex=1.1, col = "chocolate4"))
with(two, points(sesmntd ~ aspect, pch = 19, cex=0.8, col= "darkorange"))
with(one, points(sesmntd ~ aspect, pch = 19, cex=0.8, col= "gold"))

axis(1)
axis(2)
legend("bottomright", expression(~ R^2 ~ "= 0.61, p < 0.001"), text.col= "chocolate4", box.lty=0, inset=0.04, cex=0.8)
mtext("(a)", cex = 0.8, side = 3, adj = -0.23, line = 0.1)

# PD ~ aspect
pdCIa <- visregList(visreg(pd3.a, plot=FALSE), collapse=TRUE)
plot(pdCIa, overlay=TRUE, line=list(lwd=0),
     fill=list(col=c("#D55E006E")),
     points=list(cex=0), xlab = "Aspect, S = -1, N = 1", ylab="Faith's PD", legend=FALSE, cex.lab=1.1, ylim=c(500,2000), axes=FALSE)
abline(pd3.a, col = "chocolate4", lwd=3)

with(data, points(pd ~ aspect, pch = 19, cex=1.1, col= "chocolate4"))
with(two, points(pd ~ aspect, pch = 19, cex=0.8, col= "darkorange"))
with(one, points(pd ~ aspect, pch = 19, cex=0.8, col= "gold"))

axis(1)
axis(2)
legend("topright", expression(~ R^2 ~ "= 0.22, p = 0.013"), text.col = "chocolate4", box.lty=0, inset=0.03, cex=0.8)
mtext("(b)", cex = 0.8, side = 3, adj = -0.23, line = 0.1)

# SESmpd ~ tpi
mpdCIt <- visregList(visreg(mpd3.tpi, plot=FALSE), collapse=TRUE)
plot(mpdCIt, overlay=TRUE, line=list(lwd=0),
     fill=list(col=c("#D55E006E")),
     points=list(cex=0), xlab = "Terrain Position Index", ylab=expression(SES[MPD]), legend=FALSE, cex.lab=1.1, xlim=c(-1,1), ylim=c(-3,3), axes=FALSE)
abline(mpd3.tpi, col = "chocolate4", lwd=3)

with(data, points(sesmpd ~ tpi, pch = 19, cex=1.1, col= "chocolate4"))
with(two, points(sesmpd ~ tpi, pch = 19, cex=0.8, col= "darkorange"))
with(one, points(sesmpd ~ tpi, pch = 19, cex=0.8, col= "gold"))

axis(1)
axis(2)
legend("bottomright", expression(~ R^2 ~ "= 0.20, p = 0.018"), text.col = "chocolate4", box.lty=0, inset=0.03, cex=0.8)
mtext("(c)", cex = 0.8, side = 3, adj = -0.23, line = 0.1)

dev.off()
```
