## Appendix 2 for "Fractal triads efficiently sample ecological diversity and processes across spatial scales": comm-diversity-across-env-and-scale-fig3.R

```
# Assessing community (phylogenetic) diversity across environment - 2018 data - Bayesian method
# Elizabeth Simpson & Will Pearse - 2017-9-21
#setwd("~/Documents/projects/fractal_sampling_div_rhf/analysis_code") # uncomment to set working directory on computer

###########
# HEADERS #
###########
library(pez)
library(vegan)
library(lme4)
library(visreg) # need to load this package to plot loaded .Rdata

# functions for splitting the data into different triad levels
subset.first.level <- function(data){
  one <- matrix(nrow=0,ncol=ncol(data))
  colnames(one) <- colnames(data)
  for (i in seq_len(nrow(data))){
    if (data$first[i] == data$second[i] & data$second[i] == data$third[i] & data$third[i] == data$fourth[i]){
      one <- rbind(one, data[i,])
    }
  }
  return(one)
}

subset.second.level <- function(data){
  two <- matrix(nrow=0,ncol=ncol(data))
  colnames(two) <- colnames(data)
  for (i in seq_len(nrow(data))){
    if (data$second[i] == data$third[i] & data$third[i] == data$fourth[i]){
      two <- rbind(two, data[i,])
    }
  }
  return(two)
}  

subset.third.level <- function(data){
  three <- matrix(nrow=0,ncol=ncol(data))
  colnames(three) <- colnames(data)
  for (i in seq_len(nrow(data))){
    if (data$third[i] == data$fourth[i]){
      three <- rbind(three, data[i,])
    }
  }
  return(three)
}

r2_and_p <- function(model){
  return(c(round(summary(model)$r.squared,2), round(pf(summary(model)$fstatistic[1],summary(model)$fstatistic[2],summary(model)$fstatistic[3],lower.tail=FALSE),3)))
}

##############################################################################
# Make comparative community object from raw precense & percent cover data ###
##############################################################################

################
# Community data 

# load in cover data, reformat, remove NAs, remove species essentially there as notes
data <- read.csv("../data/rhf_2018_cover.csv", as.is=TRUE)
comm <- with(data, tapply(Cover, list(Plot_id,Species), function(x) mean(x,na.rm=TRUE)))
comm[is.na(comm)] <- 0
comm <- comm[,!grepl("^[a-z]+", colnames(comm))]
comm <- comm[,!grepl("\\(|/", colnames(comm))]
comm <- comm[,!grepl("sp\\.", colnames(comm))]
colnames(comm) <- tolower(colnames(comm))

################################
# Phylogeny (Zanne et. al. 2014) 

# load tree, reformat sp names, create combined data object
tree <- read.tree("../data/Vascular_Plants_rooted.dated.tre")
tree$tip.label <- tolower(gsub("_", " ", tree$tip.label))
tree <- congeneric.merge(tree, colnames(comm), split = " ")

####################
# Environmental data

# load in terrain variables, name rows with plot ID's, create comparative community object
env <- read.csv("../data/clean_rhf_terrain.csv", as.is=TRUE)
rownames(env) <- env$plot_id
c.data <- comparative.comm(tree, comm, env=env)

###################################
### Calculate diversity metrics ###
###################################

# Calculte divresiy metrics from abundance community data - 
ses.mpd <- .ses.mpd(c.data) 
ses.mntd <- .ses.mntd(c.data)
pd <- .pd(c.data)
simps.div <- diversity(c.data$comm, index="simpson")

# Compile metrics into a data frame with environmental information
data <- cbind(c.data$env, ses.mpd$ntaxa, ses.mpd$mpd.obs.z, ses.mntd$mntd.obs.z, pd[,1], simps.div)
data <- data[,c(3:15)]
colnames(data) <- c("first", "second", "third", "fourth", "elev", "aspect", "slope", "tpi", "ntaxa", "sesmpd", "sesmntd", "pd", "simpsdiv")

# Subset the data into different triad levels
one <- subset.first.level(data)
two <- subset.second.level(data)
three <- subset.third.level(data)
# four is the whole data frame of all 78 surveyed plots

#########################################
# Modeling diversity across environment #

# Species richness
nsp4.all <- with(data, lm(ntaxa ~ elev + slope + aspect + tpi)) # no sig.
nsp <- with(data, lm(ntaxa ~ 1)) 

# SES_mpd (z) 
mpd4.all <- with(data, lm(sesmpd ~ elev + slope + aspect + tpi)) # elev + aspect sig.

mpd4.e <- with(data, lm(sesmpd ~ elev))
mpd4.a <- with(data, lm(sesmpd ~ aspect))
mpd3.a <- with(three, lm(sesmpd ~ aspect))

# SES_mntd (z) 
mntd4.all <- with(data, lm(sesmntd ~ elev + slope + aspect + tpi)) # elev + aspect sig.

mntd <- with(data, lm(sesmntd ~ 1))

mntd4.a <- with(data, lm(sesmntd ~ aspect))
mntd3.a <- with(three, lm(sesmntd ~ aspect))
mntd2.a <- with(two, lm(sesmntd ~ aspect))

# Faith's PD
pd4.all <- with(data, lm(pd ~ elev + slope + aspect + tpi)) # aspect sig.

pd <- with(data, lm(pd ~ 1))

pd4.a <- with(data, lm(pd ~ aspect))
pd3.a <- with(three, lm(pd ~ aspect))

# Simpson's Div ####
simps4.all <- with(data, lm(simpsdiv ~ elev + slope + aspect + tpi)) # no sig
simps <- with(data, lm(simpsdiv ~ 1)) # value is 0.71

# output
output <- rbind(r2_and_p(mpd4.e), r2_and_p(mpd4.a), r2_and_p(mpd3.a), r2_and_p(mntd4.a), r2_and_p(mntd3.a), r2_and_p(mntd2.a), r2_and_p(pd4.a), r2_and_p(pd3.a))
rownames(output) <- c("mpd4.e", "mpd4.a", "mpd3.a", "mntd4.a", "mntd3.a", "mntd2.a", "pd4.a", "pd3.a")
colnames(output) <- c("r_squared", "p_value")

write.csv(output, "./figures-results/comm-2018-results.csv")

##################################################################################################################################################
# Plot Figure 3 ##############################################################################################################################
#setwd("~/Documents/projects/fractal_sampling_div_rhf/analysis_code/figures")
jpeg("./figures-results/fbsd-fig3.jpeg", width = 9, height = 8, units = 'in', res = 300) # Make plot
par(mfrow=c(2,2))
par(mar=c(4,4.5,3,1))
par(oma=c(.7,.7,.7,.7))

# SESmntd ~ aspect
mntdCIa <- visregList(visreg(mntd2.a, plot=FALSE), visreg(mntd3.a, plot=FALSE), visreg(mntd4.a, plot=FALSE), collapse=TRUE)
plot(mntdCIa, overlay=TRUE, line=list(lwd=0),
     fill=list(col=c("#E69F005A","#D55E006E","#0072B250")),
     points=list(cex=0), xlab= "Aspect, S = -1, N = 1", ylab=expression(SES[MNTD]), legend=FALSE, cex.lab=1.1, ylim=c(-3.2, 3), axes=FALSE)
abline(mntd2.a, col = "darkorange", lwd=3)
abline(mntd3.a, col= "chocolate4", lwd=3)
abline(mntd4.a, col= "dodgerblue4", lwd=3)

with(data, points(sesmntd ~ aspect, pch=19, cex=1.1, col = "dodgerblue4"))
with(three, points(sesmntd ~ aspect, pch = 19, cex=0.8, col= "chocolate4"))
with(two, points(sesmntd ~ aspect, pch = 19, cex=0.8, col= "darkorange"))
with(one, points(sesmntd ~ aspect, pch = 19, cex=0.8, col= "gold"))

axis(1)
axis(2)
legend("bottomright", c(expression(~ R^2 ~ "= 0.48, p < 0.001"), expression(~ R^2 ~ "= 0.51, p < 0.001"), expression(~ R^2 ~ "= 0.57, p = 0.018")), text.col=c("dodgerblue4", "chocolate4", "darkorange"), box.lty=0, inset=0.04, cex=0.8)
mtext("(a)", cex = 1, side = 3, adj = -0.18, line = 0.1)

# PD ~ aspect
pdCIa <- visregList(visreg(pd3.a, plot=FALSE), visreg(pd4.a, plot=FALSE), collapse=TRUE)
plot(pdCIa, overlay=TRUE, line=list(lwd=0),
     fill=list(col=c("#D55E006E", "#0072B250")),
     points=list(cex=0), xlab = "Aspect, S = -1, N = 1", ylab="Faith's PD", legend=FALSE, cex.lab=1.1, ylim=c(500,2000), axes=FALSE)
abline(pd3.a, col = "chocolate4", lwd=3)
abline(pd4.a, col= "dodgerblue4", lwd=3)

with(data, points(pd ~ aspect, pch=19, cex=1.1, col = "dodgerblue4"))
with(three, points(pd ~ aspect, pch = 19, cex=0.8, col= "chocolate4"))
with(two, points(pd ~ aspect, pch = 19, cex=0.8, col= "darkorange"))
with(one, points(pd ~ aspect, pch = 19, cex=0.8, col= "gold"))

axis(1)
axis(2)
legend("bottomright", c(expression(~ R^2 ~ "= 0.34, p < 0.001"), expression(~ R^2 ~ "= 0.27, p = 0.006")), text.col = c("dodgerblue4", "chocolate4"), box.lty=0, inset=0.03, cex=0.8)
mtext("(b)", cex = 1, side = 3, adj = -0.18, line = 0.1)

# SESmpd ~ aspect
mpdCIa <- visregList(visreg(mpd3.a, plot=FALSE), visreg(mpd4.a, plot=FALSE), collapse=TRUE)
plot(mpdCIa, overlay=TRUE, line=list(lwd=0),
     fill=list(col=c("#D55E006E", "#0072B250")),
     points=list(cex=0), xlab="Aspect, S = -1, N = 1", ylab=expression(SES[MPD]), legend=FALSE, cex.lab=1.1, ylim=c(-3,2), axes=FALSE)
abline(mpd3.a, col= "chocolate4", lwd=3) 
abline(mpd4.a, col= "dodgerblue4", lwd=3)

with(data, points(sesmpd ~ aspect, pch=19, cex=1.1, col = "dodgerblue4"))
with(three, points(sesmpd ~ aspect, pch = 19, cex=0.8, col= "chocolate4"))
with(two, points(sesmpd ~ aspect, pch = 19, cex=0.8, col= "darkorange"))
with(one, points(sesmpd ~ aspect, pch = 19, cex=0.8, col= "gold"))

axis(1)
axis(2)
legend("bottomright", c(expression(~ R^2 ~ "= 0.12, p = 0.002"), expression(~ R^2 ~ "= 0.18, p = 0.030")), text.col=c("dodgerblue4", "chocolate4"), box.lty=0, inset=0.03, cex=0.8)
mtext("(c)", cex = 1, side = 3, adj = -0.18, line = 0.1)

# SESmpd ~ elevation
mpdCIe <- visregList(visreg(mpd4.e, plot=FALSE), collapse=TRUE)
plot(mpdCIe, overlay=TRUE, line=list(lwd=0),
     fill=list(col=c("#0072B250")),
     points=list(cex=0), xlab= "Elevation (m)", ylab = expression(SES[MPD]), legend=FALSE, cex.lab=1.1, ylim=c(-3,2), axes=FALSE)
abline(mpd4.e, col= "dodgerblue4", lwd=3)

with(data, points(sesmpd ~ elev, pch=19, cex=1.1, col = "dodgerblue4"))
with(three, points(sesmpd ~ elev, pch = 19, cex=0.8, col= "chocolate4"))
with(two, points(sesmpd ~ elev, pch = 19, cex=0.8, col= "darkorange"))
with(one, points(sesmpd ~ elev, pch = 19, cex=0.8, col= "gold"))

axis(1)
axis(2)
legend("bottomright", expression(~ R^2 ~ "= 0.10, p = 0.005"), text.col="dodgerblue4", box.lty=0, inset=0.03, cex=0.8)
legend("topright", cex=0.8, c("fourth", "third", "second", "first"), col=c("dodgerblue4","chocolate4","darkorange","gold"), lty=1, lwd=4, pch=19, inset=0.03)
mtext("(d)", cex = 1, side = 3, adj = -0.18, line = 0.1)
dev.off()

#################################################################################################################################################
# Plot Figure 3 supplement ##################################################################################################################
jpeg("./figures-results/fbsd-fig3-supplement.jpeg", width = 8.5, height = 11, units = 'in', res = 300)
par(mfrow=c(5,2))
par(mar=c(3.8,4,0,0))
par(oma=c(1,1,2,2))

# SESmntd ~ elevation
with(data, plot(sesmntd ~ elev, pch=19, cex=1.1, col = "dodgerblue4", xlab="", ylab = expression(SES[MNTD]), cex.lab=1.1, ylim=c(-3.2,3), axes=FALSE))
abline(mntd, col="dodgerblue4", lwd=3)
with(three, points(sesmntd ~ elev, pch = 19, cex=0.8, col= "chocolate4"))
with(two, points(sesmntd ~ elev, pch = 19, cex=0.8, col= "darkorange"))
with(one, points(sesmntd ~ elev, pch = 19, cex=0.8, col= "gold"))

axis(1)
axis(2)
mtext("(a)", cex = 1 , side = 3, adj = -0.15, line = 0.1)

# SESmntd ~ aspect
mntdCIa <- visregList(visreg(mntd2.a, plot=FALSE), visreg(mntd3.a, plot=FALSE), visreg(mntd4.a, plot=FALSE), collapse=TRUE)
plot(mntdCIa, overlay=TRUE, line=list(lwd=0),
     fill=list(col=c("#E69F005A","#D55E006E","#0072B250")),
     points=list(cex=0), xlab= "Aspect, S = -1, N = 1", ylab=expression(SES[MNTD]), legend=FALSE, cex.lab=1.1, ylim=c(-3.2, 3), axes=FALSE)
abline(mntd2.a, col = "darkorange", lwd=3)
abline(mntd3.a, col= "chocolate4", lwd=3)
abline(mntd4.a, col= "dodgerblue4", lwd=3)

with(data, points(sesmntd ~ aspect, pch=19, cex=1.1, col = "dodgerblue4"))
with(three, points(sesmntd ~ aspect, pch = 19, cex=0.8, col= "chocolate4"))
with(two, points(sesmntd ~ aspect, pch = 19, cex=0.8, col= "darkorange"))
with(one, points(sesmntd ~ aspect, pch = 19, cex=0.8, col= "gold"))

axis(1)
axis(2)
legend("bottomright", c(expression(~ R^2 ~ "= 0.48, p < 0.001"), expression(~ R^2 ~ "= 0.51, p < 0.001"), expression(~ R^2 ~ "= 0.57, p = 0.018")), text.col=c("dodgerblue4", "chocolate4", "darkorange"), box.lty=0, inset=0.04, cex=0.8)
mtext("(b)", cex = 1, side = 3, adj = -0.15, line = 0.1)

# SESmpd ~ elevation
mpdCIe <- visregList(visreg(mpd4.e, plot=FALSE), collapse=TRUE)
plot(mpdCIe, overlay=TRUE, line=list(lwd=0),
     fill=list(col=c("#0072B250")),
     points=list(cex=0), xlab= "Elevation (m)", ylab = expression(SES[MPD]), legend=FALSE, cex.lab=1.1, ylim=c(-3,2), axes=FALSE)
abline(mpd4.e, col= "dodgerblue4", lwd=3)

with(data, points(sesmpd ~ elev, pch=19, cex=1.1, col = "dodgerblue4"))
with(three, points(sesmpd ~ elev, pch = 19, cex=0.8, col= "chocolate4"))
with(two, points(sesmpd ~ elev, pch = 19, cex=0.8, col= "darkorange"))
with(one, points(sesmpd ~ elev, pch = 19, cex=0.8, col= "gold"))

axis(1)
axis(2)
legend("bottomright", expression(~ R^2 ~ "= 0.10, p = 0.005"), text.col="dodgerblue4", box.lty=0, inset=0.03, cex=0.8)
mtext("(c)", cex = 1, side = 3, adj = -0.18, line = 0.1)

# SESmpd ~ aspect
mpdCIa <- visregList(visreg(mpd3.a, plot=FALSE), visreg(mpd4.a, plot=FALSE), collapse=TRUE)
plot(mpdCIa, overlay=TRUE, line=list(lwd=0),
     fill=list(col=c("#D55E006E", "#0072B250")),
     points=list(cex=0), xlab="Aspect, S = -1, N = 1", ylab=expression(SES[MPD]), legend=FALSE, cex.lab=1.1, ylim=c(-3,2), axes=FALSE)
abline(mpd3.a, col= "chocolate4", lwd=3) 
abline(mpd4.a, col= "dodgerblue4", lwd=3)

with(data, points(sesmpd ~ aspect, pch=19, cex=1.1, col = "dodgerblue4"))
with(three, points(sesmpd ~ aspect, pch = 19, cex=0.8, col= "chocolate4"))
with(two, points(sesmpd ~ aspect, pch = 19, cex=0.8, col= "darkorange"))
with(one, points(sesmpd ~ aspect, pch = 19, cex=0.8, col= "gold"))

axis(1)
axis(2)
legend("bottomright", c(expression(~ R^2 ~ "= 0.12, p = 0.002"), expression(~ R^2 ~ "= 0.18, p = 0.030")), text.col=c("dodgerblue4", "chocolate4"), box.lty=0, inset=0.03, cex=0.8)
mtext("(d)", cex = 1, side = 3, adj = -0.15, line = 0.1)

# PD ~ elevation
with(data, plot(pd ~ elev, pch=19, cex=1.1, col = "dodgerblue4", xlab= "", ylab = "Faith's PD", cex.lab=1.1, ylim=c(500, 2000), axes=FALSE))
abline(pd, col= "dodgerblue4", lwd=3)
with(three, points(pd ~ elev, pch = 19, cex=0.8, col= "chocolate4"))
with(two, points(pd ~ elev, pch = 19, cex=0.8, col= "darkorange"))
with(one, points(pd ~ elev, pch = 19, cex=0.8, col= "gold"))

axis(1)
axis(2)
mtext("(e)", cex = 1, side = 3, adj = -0.15, line = 0.1)

# PD ~ aspect
pdCIa <- visregList(visreg(pd3.a, plot=FALSE), visreg(pd4.a, plot=FALSE), collapse=TRUE)
plot(pdCIa, overlay=TRUE, line=list(lwd=0),
     fill=list(col=c("#D55E006E", "#0072B250")),
     points=list(cex=0), xlab = "Aspect, S = -1, N = 1", ylab="Faith's PD", legend=FALSE, cex.lab=1.1, ylim=c(500,2000), axes=FALSE)
abline(pd3.a, col = "chocolate4", lwd=3)
abline(pd4.a, col= "dodgerblue4", lwd=3)

with(data, points(pd ~ aspect, pch=19, cex=1.1, col = "dodgerblue4"))
with(three, points(pd ~ aspect, pch = 19, cex=0.8, col= "chocolate4"))
with(two, points(pd ~ aspect, pch = 19, cex=0.8, col= "darkorange"))
with(one, points(pd ~ aspect, pch = 19, cex=0.8, col= "gold"))

axis(1)
axis(2)
legend("bottomright", c(expression(~ R^2 ~ "= 0.34, p < 0.001"), expression(~ R^2 ~ "= 0.27, p = 0.006")), text.col = c("dodgerblue4", "chocolate4"), box.lty=0, inset=0.03, cex=0.8)
mtext("(f)", cex = 1, side = 3, adj = -0.15, line = 0.1)

# nsp ~ elevation
with(data, plot(ntaxa~ elev, pch=19, cex=1.1, col = "dodgerblue4", xlab = "", ylab = "Species Richness", cex.lab=1.1, ylim=c(0,20), axes=FALSE))
abline(nsp, col= "dodgerblue4", lwd=3)
with(three, points(ntaxa ~ elev, pch = 19, cex=0.8, col= "chocolate4"))
with(two, points(ntaxa ~ elev, pch = 19, cex=0.8, col= "darkorange"))
with(one, points(ntaxa ~ elev, pch = 19, cex=0.8, col= "gold"))

axis(1)
axis(2)
mtext("(g)", cex = 1, side = 3, adj = -0.15, line = 0.1)

# nsp ~ aspect
with(data, plot(ntaxa ~ aspect, pch=19, cex=1.1, col = "dodgerblue4", xlab = "", ylab = "Species Richness", cex.lab=1.1, ylim=c(0,20), axes=FALSE))
abline(nsp, col= "dodgerblue4", lwd=3)
with(three, points(ntaxa ~ aspect, pch = 19, cex=0.8, col= "chocolate4"))
with(two, points(ntaxa ~ aspect, pch = 19, cex=0.8, col= "darkorange"))
with(one, points(ntaxa ~ aspect, pch = 19, cex=0.8, col= "gold"))

axis(1)
axis(2)
mtext("(h)", cex = 1, side = 3, adj = -0.15, line = 0.1)

# simpsdiv ~ elevation
with(data, plot(simpsdiv ~ elev, pch=19, cex=1.1, col = "dodgerblue4", xlab = "Elevation", ylab = "Simpson's Diversity", cex.lab=1.1, ylim=c(0.3,1), axes=FALSE))
abline(simps, col= "dodgerblue4", lwd=3)
with(three, points(simpsdiv ~ elev, pch = 19, cex=0.8, col= "chocolate4"))
with(two, points(simpsdiv ~ elev, pch = 19, cex=0.8, col= "darkorange"))
with(one, points(simpsdiv ~ elev, pch = 19, cex=0.8, col= "gold"))

axis(1)
axis(2)
mtext("(i)", cex = 1, side = 3, adj=-0.15, line = 0.1)

# simpsdiv ~ aspect
with(data, plot(simpsdiv ~ aspect, pch=19, cex=1.1, col = "dodgerblue4", xlab="Aspect, S=-1, N=1", ylab = "", cex.lab=1.1, ylim=c(.3,1), axes=FALSE))
abline(simps, col= "dodgerblue4", lwd=3)
with(three, points(simpsdiv ~ aspect, pch = 19, cex=0.8, col= "chocolate4"))
with(two, points(simpsdiv ~ aspect, pch = 19, cex=0.8, col= "darkorange"))
with(one, points(simpsdiv ~ aspect, pch = 19, cex=0.8, col= "gold"))

axis(1)
axis(2)
legend("bottom", cex=0.5, c("fourth", "third", "second", "first"), col=c("dodgerblue4","chocolate4","darkorange","gold"), lty=1, lwd=4, pch=19, inset=0.03)
mtext("(j)", cex = 1, side = 3, adj = -0.15, line = 0.1)
dev.off()
```
