## Appendix 2 for "Fractal triads efficiently sample ecological diversity and processes across spatial scales": environmental-variables-2017-supplement.R

```
# Environmental variables from DEM's and measured terrain #
# Elizabeth Simpson - 2017-10-26 ##########################
# most recently neatened 2019 - 4 - 07 ####################
setwd("~/Documents/projects/fractal_sampling_div_rhf")

###########
# Headers #
###########
library(raster) 
library (rgdal)


# cosine transform aspect (coverts from degrees to radians within function) S = -1, N = 1
cos.transform <- function(asp_deg){
  asp_cos <- NA
  for (i in 1:length(asp_deg)){
    asp_cos[i] <- cos((asp_deg[i])*pi/180)
  }
  return(asp_cos)
}

calc.slope <- function(slope_up, slope_down){
  slope_deg <- NA
  for (i in 1: length(slope_up)){
    slope_deg[i] <- mean(c(slope_up[i], slope_down[i]))
  }
  return(slope_deg)
}

#################
# Measured data #
#################

# load and clean measured terrain
m.terrain <- read.csv("./data/rhf_2018_terrain_measured.csv", as.is=TRUE)
m.terrain <- m.terrain[, c(2:6)] #relevant data
m.terrain <- m.terrain[-79,] #remove plot 1000

# transform aspect
m.terrain$m_cos_asp <- with(m.terrain, cos.transform(Aspect_deg))

# calculate slope
m.terrain$slope_deg <- with(m.terrain, calc.slope(Slope_down_deg, Slope_up_deg))

##############################
# GIS data ###################
##############################

# load raster DEM and reproject raster
dm <- raster("../DEM_5m_12TVM400200.asc")
#dm <- raster("https://storage.googleapis.com/state-of-utah-sgid-downloads/auto-correlated-dems/5-meter/12TVM400200.zip")
# I only want the file 12TVM400200.asc from this...but I can't seem to get this unzipped and loaded in R
crs(dm) <- "+proj=utm +zone=12 +ellps=GRS80 +towgs84=0,0,0,0,0,0,0 +units=m +no_defs"
shp_18 <- shapefile("./data/2018plots/2018_plot_locs.shp")
shp_17 <- shapefile("./data/2017plots/2017_plot_locs.shp")

# transform coordinates
shp_utm_18 <- spTransform(shp_18, crs(dm))
shp_utm_17 <- spTransform(shp_17, crs(dm))

# create occurences of unsampled plots
plots <- cbind(as.data.frame(shp_utm_18@data$name),as.data.frame(shp_utm_18@coords))
second <- plots[c(9,29,30,33,60,65),]
first <- plots[c(2,36,45),]
notsamp_plots <- plots[c(15,58),]

# create extent window to crop DEM to match site area
ext <- extent(shp_utm_18)
ext@xmin <- ext@xmin-100
ext@ymin <- ext@ymin-100
ext@xmax <- ext@xmax+100
ext@ymax <- ext@ymax+100

#crop DEM
dm.cr <- crop(dm, ext)

#################################
# Extract terrain vars from DEM #
#################################

# Choose which terrain vars to extract
asp <- terrain(dm.cr, opt='aspect', unit='degrees', neighbors=8)
slp <- terrain(dm.cr, opt='slope', unit='degrees', neighbors=8)
tpi <- terrain(dm.cr, opt=c('TPI'))

dem.terrain <- data.frame(shp_utm_18$name, 
                          extract(dm, shp_utm_18),
                          extract(asp, shp_utm_18),
                          extract(slp, shp_utm_18),
                          extract(tpi, shp_utm_18))

# name the columns something sensible
colnames(dem.terrain) <- c("plot_id","dem_elev","dem_asp", "dem_slope","TPI")

# pull out info from 2322
terrain2322 <- subset(dem.terrain, (plot_id == 2322))
terrain2322$m_cos_asp <- with(terrain2322, cos.transform(dem_asp))
terrain2322 <- terrain2322[,c("plot_id","dem_elev","m_cos_asp","dem_slope", "TPI")]
colnames(terrain2322) <- c("plot_ID","elev","aspect","slope", "TPI")

# remove plots not surveyed
dem.terrain <- subset(dem.terrain, !(plot_id == 2312))
dem.terrain <- subset(dem.terrain, !(plot_id == 2321))
dem.terrain <- subset(dem.terrain, !(plot_id == 2322))

# create data from of all env/terrain variables
terrain <- merge(m.terrain, dem.terrain, by.x="Plot_id", by.y="plot_id")

# clean
terrain <- terrain[,c("Plot_id", "gps_elev_m", "m_cos_asp", "slope_deg","TPI")]
colnames(terrain) <- c("plot_ID", "elev", "aspect", "slope", "TPI")
terrain <- rbind(terrain, terrain2322)

# separate plot_id - for spatially exlicit analysis
library(tidyverse) #can't put in header bc extract() function conflicts with extract() in raster package
terrain <- terrain %>% extract(plot_ID, c("first", "second", "third", "fourth"), "([[:digit:]]+)([[:digit:]]+)([[:digit:]]+)([[:digit:]]+)", remove = FALSE)

# export clean terrain file
write.csv(terrain, file="./data/clean_rhf_terrain_2017.csv")

#############################
# Plot plots ################
#############################
setwd("~/Documents/projects/fractal_sampling_div_rhf/analysis_code/figures")
jpeg("fbsd-fig1-raw.jpeg", width = 9, height = 8, units = 'in', res = 300)
plot(dm.cr, col=gray.colors(10, start=0.95, end=0.2), useRaster= F, xlim=c(ext@xmin,ext@xmax), ylim=c(ext@ymin,ext@ymax), xaxt = "n", yaxt="n", bty="n")
# add all 81 plots
points(shp_utm_18, pch=19, cex=1.2)
points(shp_utm_18, pch=19, cex=1, col = "#0072B2")

# add ones established and surveyed in 2017
points(shp_utm_17, pch=19, col="#D55E00FF")

# add other triad levels
points(second[,2:3], pch=19, col="#E69F00FF")
points(first[,2:3], pch=19, col="#F0E442FF")
# add ones not sampled
points(notsamp_plots[,2:3], pch=19)
dev.off()
```
