## Appendix 2 for "Fractal triads efficiently sample ecological diversity and processes across spatial scales": fractal-random-comparison-fig2.R

```
# Number of comparisons for different distance classes #####################
# Will Pearse # 2018-01-22 #################################################
# adjusted to build evenly by Elizabeth Simpson & Guen Grosklos 2018-10-12 #

###########
# HEADERS #
###########

# Build fractal sampling design 
# These functions build a fractal-based sampling design based on four arguments: 
# 1 and 2 - x and y - coordinates of the starting vertex
# 3 - triads - the number of triad levels (>=1) in the fractal design
# 4 - d - the maximum sampling distance

# From each intial vertex, make.vertices generates two additional verticies in an equilateral triangle
# This function tracks which triad level it is calulating verticies within and adjusts the distance between verticies accordingly.
make.vertices <- function(x,y,triads,d){
  size <- d/3^(triads-1)
  LV <- c(x,y)
  RV <- LV + c(size, 0)
  UV <- LV + c(size/2, (sqrt(size^2-(size/2)^2)))
  return(rbind(LV,RV,UV))
}

# From the starting vertex (LV), make.fractal uses make.verticies to add additional vertices (RV, UV) at the maximum sampling distance to make the first triad level.
# For each successive triad level, make.verticies  adds verticies at a third of the distance from the level before.
# make.fractal only adds vertices if a vertex does not already exist at that coordinate.
make.fractal <- function(x,y,triads,d){
  LV <- matrix(c(x,y),ncol=2)
  RV <- matrix(nrow=0, ncol=2)
  UV <- matrix(nrow=0, ncol=2)
  for (i in 1:triads){
    for(j in seq_len(nrow(LV))){
      tri <- make.vertices(LV[j,1],LV[j,2], i, d)
      if (i == triads){
        RV <- rbind(RV, tri[2,])
        UV <- rbind(UV, tri[3,])
      }
      if (i < triads){
        size <- d/3^(i-1)
        LV <- rbind(LV, LV[j,] + c((2/3)*size, 0))
        LV <- rbind(LV, LV[j,] + c(size/3, sqrt(((2/3)*size)^2 -(size/3)^2)))
      }
      
    }
  }
  return(rbind(LV,RV,UV))
}

# function to generate comparisons across distances sampled by fractal vs. random sampling scheme
comparison.distance <- function(triads,extent){
  output <- matrix(nrow=40, ncol=4)
  fractal <- make.fractal(x=0,y=0, triads=triads, d=extent)
  # Do randomisations
  bins <- seq(0,round(sqrt(2000^2 + 2000^2), -3),by=50)
  counts <- matrix(nrow=1000, ncol=length(bins)-1)
  for(i in seq_len(nrow(counts))){
    rnd.points <- t(replicate(nrow(fractal),runif(2,0,max(as.numeric(fractal)))))
    counts[i,] <- as.numeric(table(cut(as.numeric(dist(rnd.points)), bins)))
  }
  medians <- apply(counts, 2, median)
  sds <- apply(counts, 2, sd)
  obs.density <- as.numeric(table(cut(as.numeric(dist(fractal)), bins)))
  centers <- bins[-1] - 25
  #trim it down
  output[,1] <- obs.density[1:40]
  output[,2] <- centers[1:40]
  output[,3] <- sds[1:40]
  output[,4] <- medians[1:40]
  colnames(output) <- c("obs.density", "centers", "sds", "medians")
  return(output)
}

##########################################################################################################3
# calculate comparison across distance
two.triads <- as.data.frame(comparison.distance(2,1900))
three.triads <- as.data.frame(comparison.distance(3, 1900))
four.triads <- as.data.frame(comparison.distance(4, 1900))

# plotting
setwd("~/Documents/projects/fractal_sampling_div_rhf/analysis_code/figures")
jpeg("fbsd-fig2-raw.jpeg", width = 12, height = 4, units = 'in', res = 300)
par(mfrow=c(1,3))

# two triads
with(two.triads, plot(obs.density ~ centers, col=ifelse(obs.density > medians, "red", "blue"), ylim=c(0,12), xlab="", ylab = "", bty="l", type="n", axes=FALSE))
axis(1, cex.axis=2)
axis(2, cex.axis=2)
with(two.triads, points(medians ~ centers, col="grey40", pch=19, cex=2))
with(two.triads, arrows(centers, medians-sds, centers, medians+sds, col="grey40", code=3, angle=90, length = 0.0, lwd = 3))
with(two.triads, points(obs.density ~ centers, col=ifelse(obs.density > medians, "red", "blue"), pch=19, cex=2))

# three triads
with(three.triads, plot(obs.density ~ centers, col=ifelse(obs.density > medians, "red", "blue"), ylim=c(0,40), xlab="", ylab = "", bty="l", type="n", axes=FALSE))
axis(1, cex.axis=2)
axis(2, cex.axis=2)
with(three.triads, points(medians ~ centers, col="grey40", pch=19, cex=2))
with(three.triads, arrows(centers, medians-sds, centers, medians+sds, col="grey40", code=3, angle=90, length = 0.0, lwd = 3))
with(three.triads, points(obs.density ~ centers, col=ifelse(obs.density > medians, "red", "blue"), pch=19, cex=2))

# plotting 4 triads comparisons
with(four.triads, plot(obs.density ~ centers, col=ifelse(obs.density > medians, "red", "blue"), ylim=c(0,250), xlab="", ylab = "", bty="l", type="n", axes=FALSE))
axis(1, cex.axis=2)
axis(2, cex.axis=2)
with(four.triads, points(medians ~ centers, col="grey40", pch=19, cex=2))
with(four.triads, arrows(centers, medians-sds, centers, medians+sds, col="grey40", code=3, angle=90, length = 0.0, lwd = 3))
with(four.triads, points(obs.density ~ centers, col=ifelse(obs.density > medians, "red", "blue"), pch=19, cex=2))
dev.off()
```
