## Appendix 2 for "Fractal triads efficiently sample ecological diversity and processes across spatial scales": variance-components-analysis-2017-supplement.R

```
# Variance Components Analysis - 2017 data
# Elizabeth Simpson & Will Pearse - 2017-11-6
# Page 691 in The R Book: Mixed effects models plus Bayesian methods in rstanarm
#setwd("~/Documents/projects/fractal_sampling_div_rhf/analysis_code")
#load("./figures-results/comm-vca-2017.Rdata")

############
# Headers #
###########
library(vegan)
library(tidyverse)
library(rstanarm)
library(parallel)
library(vioplot) # need to load this package if you want to plot from loaded .Rdata
#options(mc.cores = 12) # uncomment to run Bayesian analysis
#boot_runs <- 999 # uncomment to run Bayesian analysis

# p-value for obs < null (for obs > null use 1-"p-value")
p_value <- function(distribution, obs_point){
  return(rank(c(obs_point,distribution))[1] / (length(distribution)+1))
}

# Calculate variance composition for measured metric values - report these as a percentage of total variance
var.comp.2 <- function(metric, first, second, data, adapt_delta){
  model <- stan_lmer(metric ~ 1|first/second, data = data,
  		      iter = 5000, warmup = 4000,
  		      prior = normal(),
  		      prior_intercept = normal(),
  		      prior_aux = exponential(),
  		      prior_covariance = decov(),
  		      prior_PD = FALSE, 
  		      adapt_delta = adapt_delta)
  vars <- as.data.frame(VarCorr(model))$vcov
  names(vars) <- c(names(VarCorr(model)),"residual")
  return((vars / sum(vars)) * 100)
}

# Calculate mean and 95% confidence interval for random sample of diversity metrics measured *without* replacement 
# runs - number of times to sample
# first, second - triad levels for 3 level fractal sampling design
rnd.var.boot.2 <- function(div.met, runs, first, second, data, adapt_delta){
  mat <- matrix(NA, runs, length(div.met))
  for (i in seq_len(nrow(mat))){
    mat[i,] <-sample(div.met, length(div.met))
  }
  var <- matrix(NA, nrow(mat), 3)
  var <- mcMap(
  	function(i) var.comp.2(mat[i,], first, second, data, adapt_delta), 
  	seq_len(nrow(mat)))
  var <- do.call(rbind, var)
  colnames(var) <- c("2nd:1st", "1st","residual")
  return(var)
}

# Load data by running comm-diversity-across-env-and-space-2017-supplement.R
source("comm-diversity-across-env-and-scale-2017-supplement.R")

###################################################################################################
# Variance componenets analysis for each diversity metric, randomized version of diversity metric #

# Species richness
var_nsp <- with(data, var.comp.2(ntaxa, first, second, data, 0.999))
nsp_boot <- with(data, rnd.var.boot.2(ntaxa, boot_runs, first, second, data=data, adapt_delta=0.999))

nsp_p <- (NA)
for(i in 1:3) nsp_p[i] <- p_value(nsp_boot[,i], var_nsp[i])

# SESmpd
var_mpd <- with(data, var.comp.2(sesmpd, first, second, data, 0.999))
mpd_boot <- with(data, rnd.var.boot.2(sesmpd, boot_runs, first, second, data=data, adapt_delta=0.999))

mpd_p <- (NA)
for(i in 1:3) mpd_p[i] <- p_value(mpd_boot[,i], var_mpd[i])

# PD
var_pd <- with(data, var.comp.2(pd, first, second, data, 0.999))
pd_boot <- with(data, rnd.var.boot.2(pd, boot_runs, first, second, data, 0.999))

pd_p <- (NA)
for(i in 1:3) pd_p[i] <- p_value(pd_boot[,i], var_pd[i])

# SESmntd
var_mntd <- with(data, var.comp.2(sesmntd, first, second, data, 0.999))
mntd_boot <- with(data, rnd.var.boot.2(sesmntd, boot_runs, first, second, data, 0.999))

mntd_p <- (NA)
for(i in 1:3) mntd_p[i] <- p_value(mntd_boot[,i], var_mntd[i])

# Simps div
var_simps <- with(data, var.comp.2(simpsdiv, first, second, data, 0.9999))
simps_boot <- with(data, rnd.var.boot.2(simpsdiv, boot_runs, first, second, data, 0.9999))

simps_p <- (NA)
for(i in 1:3) simps_p[i] <- p_value(simps_boot[,i], var_simps[i])

output <- rbind(var_nsp, nsp_p, var_mpd, mpd_p, var_pd, pd_p, var_mntd, mntd_p, var_simps, simps_p)
write.csv(output, "./figures-results/VCA-2017-results.csv")

######################################################################################
# Plotting #
jpeg("./figures-results/fbsd-fig4-2017-supplement-raw.jpeg", width = 9, height = 12, units = 'in', res = 300)
par(mfrow=c(3,2))
par(mar=c(2,3,1,0.5))
par(oma=c(.7,.7,.7,.7))

# MNTD
vioplot(mntd_boot[,2], mntd_boot[,1], mntd_boot[,3], xaxt="n", yaxt="n", frame.plot=F, col="azure3", border="azure3", rectCol= "gray20", lineCol="gray20", colMed = "gray20", pchMed = "-", cex=3, ylim=c(0,100))
axis(1, labels=FALSE, at=c(1,2,3))
axis(2, cex.axis=1.4)
points(1:3, var_mntd[c(2,1,3)], pch=21, cex=2.5, col="red", lwd=3, bg=NA)
title(outer=FALSE, adj = 0.03, main=expression("(a)" ~ SES[MNTD]), cex.main=1.3)

# PD
vioplot(pd_boot[,2], pd_boot[,1], pd_boot[,3], xaxt="n", yaxt="n", frame.plot=F, col="azure3", border="azure3", rectCol= "gray20", lineCol="gray20", colMed = "gray20", pchMed = "-", cex=3, ylim=c(0,100))
axis(1, labels=FALSE, at=c(1,2,3))
axis(2, labels=FALSE)
points(1:3, var_pd[c(2,1,3)], pch=21, cex=2.5, col="red", lwd=3, bg=NA)
title(outer=FALSE, adj = 0.03, main="(b) Faith's PD",cex.main=1.3)

# MPD
vioplot(mpd_boot[,2], mpd_boot[,1],mpd_boot[,3] , xaxt="n", yaxt="n", frame.plot=F, col="azure3", border="azure3", rectCol= "gray20", lineCol="gray20", colMed = "gray20", pchMed = "-", cex=3, ylim=c(0,100))
axis(1, labels=FALSE, at=c(1,2,3))
axis(2, cex.axis=1.4)
points(c(1:3), var_mpd[c(2,1,3)], pch=21, cex=2.5, col="red", lwd=3, bg=NA)
title(outer=FALSE, adj = 0.03, main=expression("(c)" ~ SES[MPD]), cex.main=1.3)

# Species richness
vioplot(nsp_boot[,2], nsp_boot[,1], nsp_boot[,3], xaxt="n", yaxt="n", frame.plot=F, col="azure3", border="azure3", rectCol= "gray20", lineCol="gray20", colMed = "gray20", pchMed = "-", cex=3, ylim=c(0,100))
axis(1, labels=FALSE, at=c(1,2,3))
axis(2, cex.axis=1.4, labels=FALSE)
points(1:3, var_nsp[c(2,1,3)], pch=21, cex=2.5, col="red", lwd=3, bg=NA)
title(outer=FALSE, adj = 0.03, main="(d) Species richness",cex.main=1.3)

# Simpson's diversity
vioplot(simps_boot[,2], simps_boot[,1], simps_boot[,3], xaxt="n", yaxt="n", frame.plot=F, col="azure3", border="azure3", rectCol= "gray20", lineCol="gray20", colMed = "gray20", pchMed = "-", cex=3, ylim=c(0,100))
axis(1, labels = FALSE, at=c(1,2,3))
axis(2, cex.axis=1.4)
points(1:3, var_simps[c(2,1,3)], pch=21, cex=2.5, col="red", lwd=3, bg=NA)
title(outer=FALSE, adj = 0.03, main="(e) Simpson's diversity",cex.main=1.3)

dev.off()

save.image("./figures-results/comm-vca-2017.Rdata")
```
