## Appendix 2 for "Fractal triads efficiently sample ecological diversity and processes across spatial scales": variance-components-analysis-fig4.R

```
# Variance Components Analysis - 2018 data
# Elizabeth Simpson & Will Pearse - 2017-11-6
# Page 691 in The R Book: Mixed effects models plus Bayesian methods in rstanarm
#setwd("~/Documents/projects/fractal_sampling_div_rhf/analysis_code") # uncomment to look at Baysian analysis on computer
#load("./figures-results/comm-vca-2018.Rdata") # un-comment to load working directory on computer

############
# Headers #
###########
library(vegan)
library(tidyverse)
library(rstanarm)
library(parallel)
library(vioplot) # need to load this package if you want to plot from loaded .Rdata
#options(mc.cores = 12) # uncomment to run Bayesian analysis
#boot_runs <- 999 # uncomment to run Bayesian analysis

# p-value for obs < null (for obs > null use 1-"p-value")
p_value <- function(distribution, obs_point){
  return(rank(c(obs_point,distribution))[1] / (length(distribution)+1))
}

# Calculate variance composition for measured metric values - report these as a percentage of total variance
var.comp.3 <- function(metric, first, second, third, data, adapt_delta){
  model <- stan_lmer(metric ~ 1|first/second/third, data = data,
                     iter = 5000, warmup = 4000,
                     prior = normal(),
                     prior_intercept = normal(),
                     prior_aux = exponential(),
                     prior_covariance = decov(),
                     prior_PD = FALSE,
                     adapt_delta = adapt_delta)
  vars <- as.data.frame(VarCorr(model))$vcov
  names(vars) <- c(names(VarCorr(model)),"residual")
  return((vars / sum(vars)) * 100)
}

# Calculate mean and 95% confidence interval for random sample of diversity metrics measured *without* replacement 
# runs - number of times to sample
# first, second, third - triad levels for 4 level fractal sampling design
rnd.var.boot.3 <- function(div.met, runs, first, second, third, data, adapt_delta){
  mat <- matrix(NA, runs, length(div.met))
  for (i in seq_len(nrow(mat))){
    mat[i,] <-sample(div.met, length(div.met))
  }
  var <- matrix(NA, nrow(mat), 4)
  var <- mcMap(
    function(i) var.comp.3(mat[i,], first, second, third, data, adapt_delta), 
    seq_len(nrow(mat)))
  var <- do.call(rbind, var)
  colnames(var) <- c("3rd:(2nd:1st)", "2nd:1st", "1st","residual")
  return(var)
}

# Load applicable data by running comm-diversity-across-env-and-scale-fig3.R
source("comm-diversity-across-env-and-scale-fig3.R")

###################################################################################################
# Variance componenets analysis for each diversity metric, randomized version of diversity metric #

# Species richness
var_nsp <- with(data, var.comp.3(ntaxa, first, second, third, data, 0.99999))
nsp_boot <- with(data, rnd.var.boot.3(ntaxa, boot_runs, first, second, third, data, 0.99999))

nsp_p <- (NA)
for (i in 1:4){
  nsp_p[i] <- p_value(nsp_boot[,i], var_nsp[i])
}

# SESmpd
var_mpd <- with(data, var.comp.3(sesmpd, first, second, third, data, 0.99999))
mpd_boot <- with(data, rnd.var.boot.3(sesmpd, boot_runs, first, second,third, data, 0.99999))

mpd_p <- (NA)
for (i in 1:4){
  mpd_p[i] <- p_value(mpd_boot[,i], var_mpd[i])
}

# PD
var_pd <- with(data, var.comp.3(pd, first, second, third, data, 0.99999))
pd_boot <- with(data, rnd.var.boot.3(pd, boot_runs, first, second, third, data, 0.99999))

pd_p <- (NA)
for (i in 1:4){
  pd_p[i] <- p_value(pd_boot[,i], var_pd[i])
}

# SESmntd
var_mntd <- with(data, var.comp.3(sesmntd, first, second, third, data, 0.99999))
mntd_boot <- with(data, rnd.var.boot.3(sesmntd, boot_runs, first, second, third, data, 0.99999))

mntd_p <- (NA)
for (i in 1:4){
  mntd_p[i] <- p_value(mntd_boot[,i], var_mntd[i])
}

# Simps div
var_simps <- with(data, var.comp.3(simpsdiv, first, second, third, data, 0.99999))
simps_boot <- with(data, rnd.var.boot.3(simpsdiv, boot_runs, first, second, third, data, 0.99999))

simps_p <- (NA)
for (i in 1:4){
  simps_p[i] <- p_value(simps_boot[,i], var_simps[i])
}

output <- rbind(var_nsp, nsp_p, var_mpd, mpd_p, var_pd, pd_p, var_mntd, mntd_p, var_simps, simps_p)
write.csv(output, "./figures-results/VCA-2018-results.csv")

#####################################################################################
# PLOTTING #
jpeg("./figures-results/fbsd-fig4-raw.jpeg", width = 9, height = 7, units = 'in', res = 300)
par(mfrow=c(2,2))
par(mar=c(2,3,1,0.5))
par(oma=c(.7,.7,.7,.7))

# SESmntd
vioplot(mntd_boot[,3], mntd_boot[,2], mntd_boot[,1], mntd_boot[,4], xaxt="n", yaxt="n", frame.plot=F, col="azure3", border="azure3", rectCol= "gray20", lineCol="gray20", colMed = "gray20", pchMed = "-", cex=3, ylim=c(0,100))
axis(1, labels=FALSE)
axis(2, cex.axis=1.4)
points(c(2:4), var_mntd[c(2,1,4)], pch=16, cex=2, col="red")
points(1, var_mntd[3], pch=21, cex=2, col="red", lwd=3, bg=NA)
title(outer=FALSE, adj = 0.03, main=expression("(a)" ~ SES[MNTD]), cex.main=1.3)

# PD
vioplot(pd_boot[,3], pd_boot[,2], pd_boot[,1], pd_boot[,4], xaxt="n", yaxt="n", frame.plot=F, col="azure3", border="azure3", rectCol= "gray20", lineCol="gray20", colMed = "gray20", pchMed = "-", cex=3, ylim=c(0,100))
axis(1, labels=FALSE)
axis(2, labels=FALSE)
points(c(1:3), var_pd[c(3:1)], pch=16, cex=2, col="red")
points(4, var_pd[4], pch=21, cex=2, col="red", lwd=3, bg=NA)
title(outer=FALSE, adj = 0.03, main="(b) Faith's PD",cex.main=1.3)

# SESmpd
vioplot(mpd_boot[,3], mpd_boot[,2],mpd_boot[,1], mpd_boot[,4], xaxt="n", yaxt="n", frame.plot=F, col="azure3", border="azure3", rectCol= "gray20", lineCol="gray20", colMed = "gray20", pchMed = "-", cex=3, ylim=c(0,100))
axis(1, labels=FALSE)
axis(2, cex.axis=1.4)
points(c(2:4), var_mpd[c(2,1,4)], pch=16, cex=2, col="red")
points(c(1), var_mpd[c(3)], pch=21, cex=2, col="red", lwd=3, bg=NA)
title(outer=FALSE, adj = 0.03, main=expression("(c)" ~ SES[MPD]), cex.main=1.3)

# Species richness
vioplot(nsp_boot[,3], nsp_boot[,2], nsp_boot[,1], nsp_boot[,4], xaxt="n", yaxt="n", frame.plot=F, col="azure3", border="azure3", rectCol= "gray20", lineCol="gray20", colMed = "gray20", pchMed = "-", cex=3, ylim=c(0,100))
axis(1, labels=FALSE)
axis(2, labels=FALSE)
points(c(1,3), var_nsp[c(3,1)], pch=16, cex=2, col="red")
points(c(2,4), var_nsp[c(2,4)], pch=21, cex=2, col="red", lwd=3, bg=NA)
title(outer=FALSE, adj = 0.03, main="(d) Species richness",cex.main=1.3)

dev.off()

#######################################################
# Plotting Simpson's Diversity for supplement #########
jpeg("./figures-results/fbsd-fig4-supplement-raw.jpeg", width = 8.5, height = 7, units = 'in', res = 300)

vioplot(simps_boot[,3], simps_boot[,2], simps_boot[,1], simps_boot[,4], xaxt="n", yaxt="n", frame.plot=F, col="azure3", border="azure3", rectCol= "gray20", lineCol="gray20", colMed = "gray20", pchMed = "-", cex=3, ylim=c(0,100) )
axis(1, labels = FALSE)
axis(2, cex.axis=1.4)
points(1:4, var_simps[c(3:1,4)], pch=21, cex=2, col="red", lwd=3, bg=NA)
title(outer=FALSE, adj = 0.03, main="Simpson's diversity",cex.main=1.3)

dev.off()

save.image("./figures-results/comm-vca-2018.Rdata")
```
