## Appendix 2 for "Fractal triads efficiently sample ecological diversity and processes across spatial scales": rhf_2017_cover_metadata.txt

Date: International format (YYY-MM-DD)

Recorder1, Recorder2: Name of person or multiple columns of
    recorders. Everyone in the field party is an observer.

Site: Code number of site, following triad format. Starting with
    outter-most triad and working in from there.

Quadrant: Which quadrant the cover recording was made in (using the
    50cm x 50cm quadrat or an approximation of it). 1: top-left
    quadrant; 2: top-right quadrant; 3: bottom-left quadrant; 4:
    bottom-right quadrant

Species: Full Latin binomial of species observed. No common names or
    abbreviations - EVER. In the format "Genus species". If soil,
    rock, poo, etc., this can be written as-is.

Cover: 0-100 the cover recorded for the "Species". If 0 is recorded,
    write it down as 0!

Notes: Any notes or observation. Keep this to a minimum; just state if
    something was an unclear identification etc.
