## Appendix 2 for "Fractal triads efficiently sample ecological diversity and processes across spatial scales": rhf_2018_cover_metadata.txt

Date: International format (YYYY-MM-DD)

Recorder_1, Recorder_2, etc.: Name of person or multiple columns of
    recorders. Everyone in the field party is an observer.

Plot_id: Code number of plot (originally called 'Site' in 2017), following triad format. Starting with outer-most triad and working in from there.

Quadrant: Which quadrant the cover recording was made in (using the
    50cm x 50cm quadrat or an approximation of it). 1: top-left
    quadrant; 2: top-right quadrant; 3: bottom-left quadrant; 4:
    bottom-right quadrant

Species: Full Latin binomial of species observed. No common names or
    abbreviations - EVER. In the format "Genus species". If soil,
    rock, poo, or place-holder unidentifid morphotype name, etc., this is written as-is but ALWAYS in lowercase. Debris like leaves, sticks, bark, etc. that can be identified from a specific species are noted as such with the type of debris in parenthesis following the "Genus species" name. Potential hybrids are identified to the species they match closest to. 

Cover: 0-100 the cover recorded for the "Species". No cover is recorded as 0. 0 is also recorded for species not rooted in the plot but overhanging the plot.
