## Appendix 2 for "Fractal triads efficiently sample ecological diversity and processes across spatial scales": rhf_2018_data_log

~/Documents/projects/fractal_sampling_div_rhf

	./raw_data/
		rhf_2018_cover.csv - Cover data from 2018 - collected by Elizabeth Simpson, Will Pearse, and Emily James and IDed by Elizabeth Simpson, Mary Barkworth, and Michael Piep
		rhf_2018_cover_metadata.txt - Metadata from 2018 cover data
		Vascular_Plants_rooted.dated.tre - phylogeny from Zanne et al. 2015 used to build trees for phylogenetic analysis
		rhf_2018_terrain_measured.csv - Elevation, slope, and aspect measured in the field
		rhf_2018_terrain_measured_metadata.txt - Metadata from 2018 terrain measured
		rhf_all_plot_locs.csv - Coordinates for all plot locations
		./2017plots/ - contains all GIS shapefiles for 2017 plots
		./2018plots/ - contains all GIS shapefiles for 2018 plots
		./SWGAP-landcover/ - contains all GIS shapefiles for assessing landcover types


		


		
