## Appendix 2 for "Fractal triads efficiently sample ecological diversity and processes across spatial scales": rhf_2018_terrain_measured_metadata.txt

Measurer: Name of person who took the topographic measurements. 

Plot_id: Code number of plot (originally called 'Site' in 2017), following triad format. Starting with outer-most triad and working in from there.

Aspect_deg: Aspect of hill plot is on, measured in degrees. Taken with a compass, by facing as directly downhill as possible and aligning the North arrow and reading off the bearing

Slope_down_deg, Slope_up_deg: Slope of hill plot is on, measured in degrees. Taken with a clinometer, by measuring the degrees from horizontal that an oject downhill (and uphill) at as close to possible at eye level is at. The mean of these two will equal the measured slope.
*** Note: not measured for 1231 through 2133 because I did not initially have the clinometer and did not keep taking slope measurements after the first few plots when collecting soil in August. 

Dist: Distance, in meters, of this plot from other plots in the largest triad level it's a part of . 

Triad_level: Which (largest) triad level this plot is part of. '

N_samples: minimum number of samples possible at this plot's largest triad level

Notes: any pertinent information about the measured terrain at this plot location
