## Supplementary figures and images for "Fractal triads efficiently sample ecological diversity and processes across spatial scales"

### fbsd-fig1-raw.jpeg

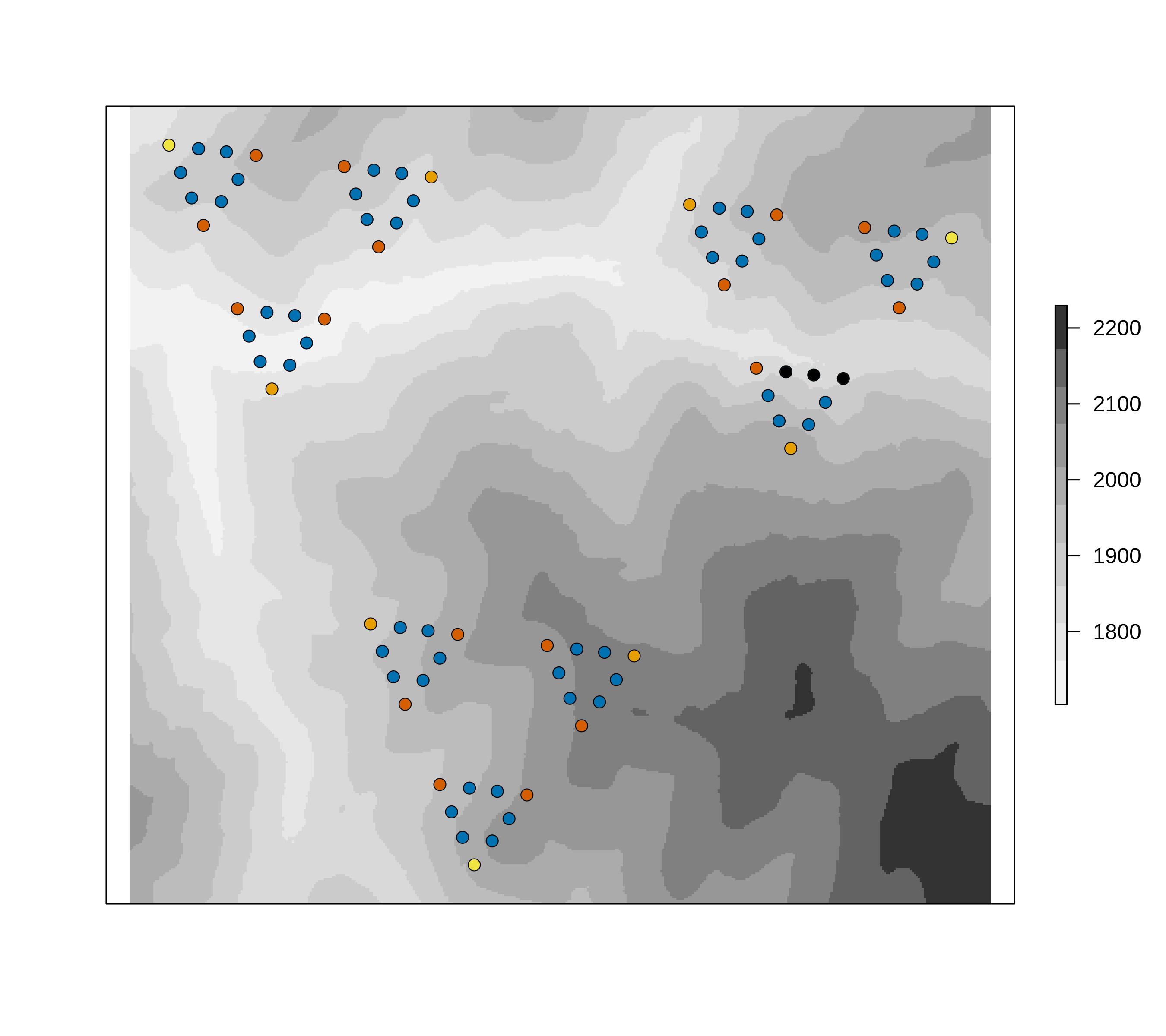

### fbsd-fig1.jpeg

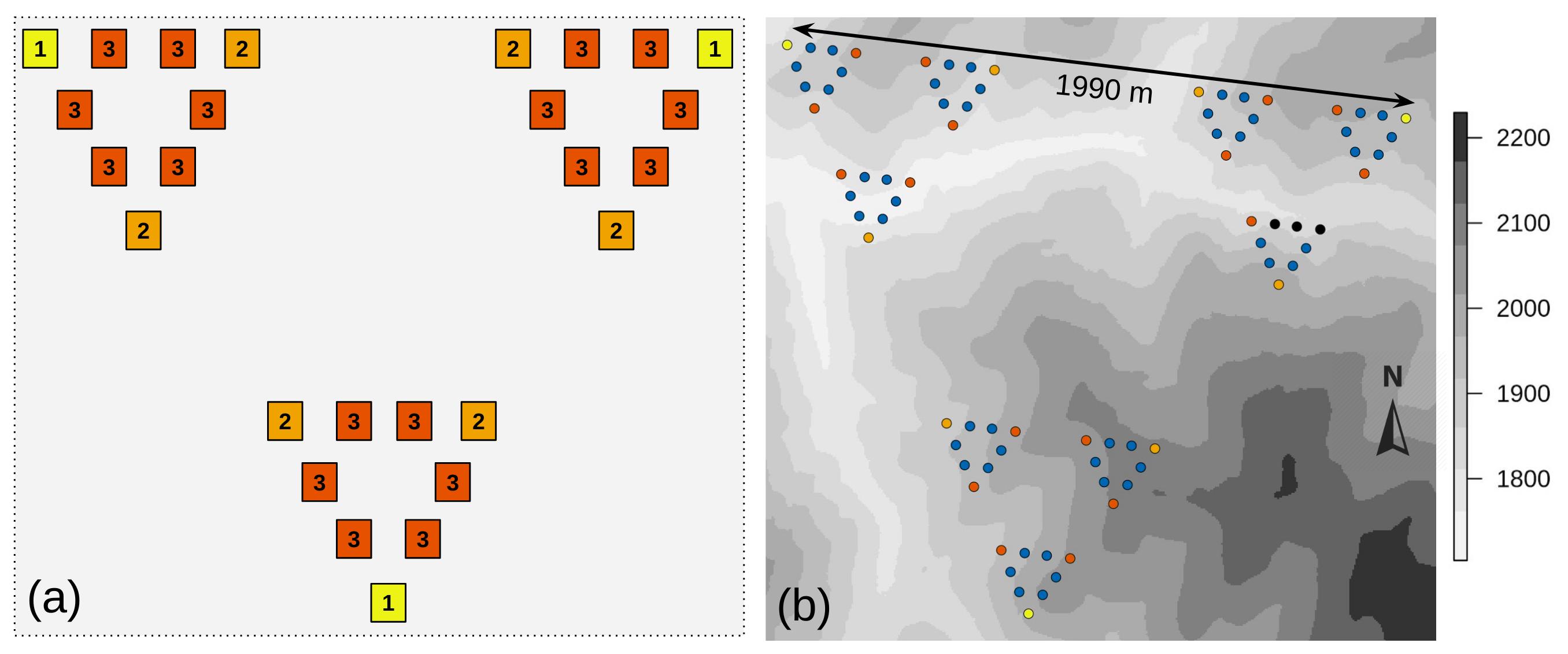

### fbsd-fig2-raw.jpeg

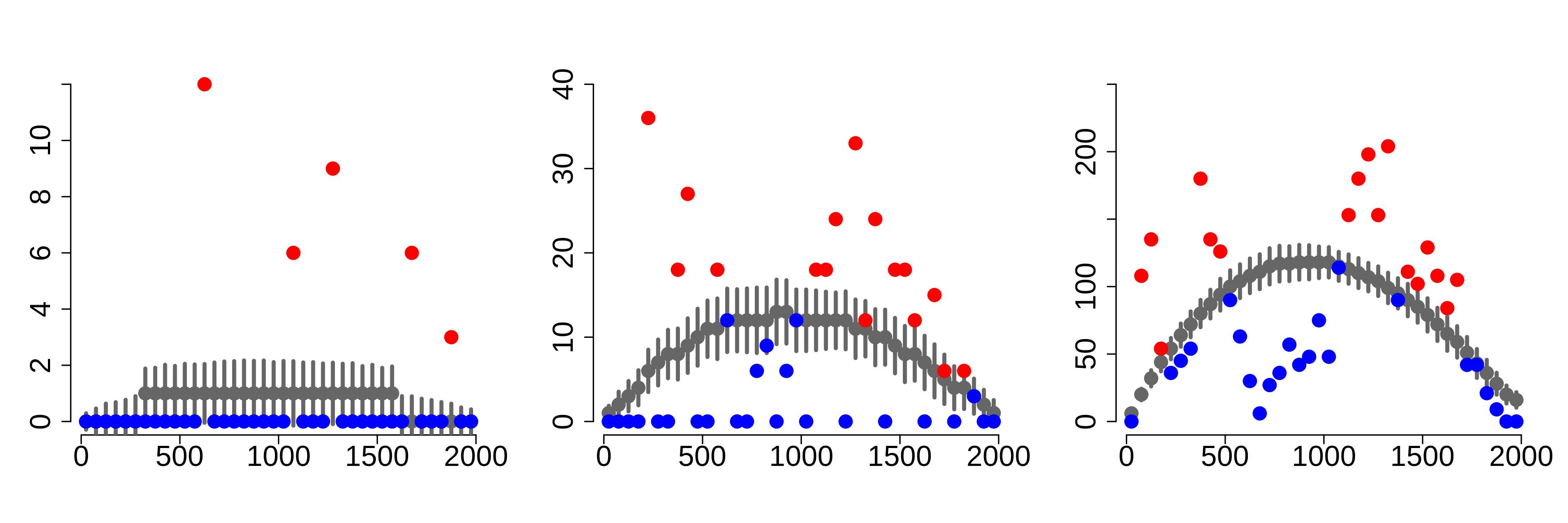

### fbsd-fig2.jpeg

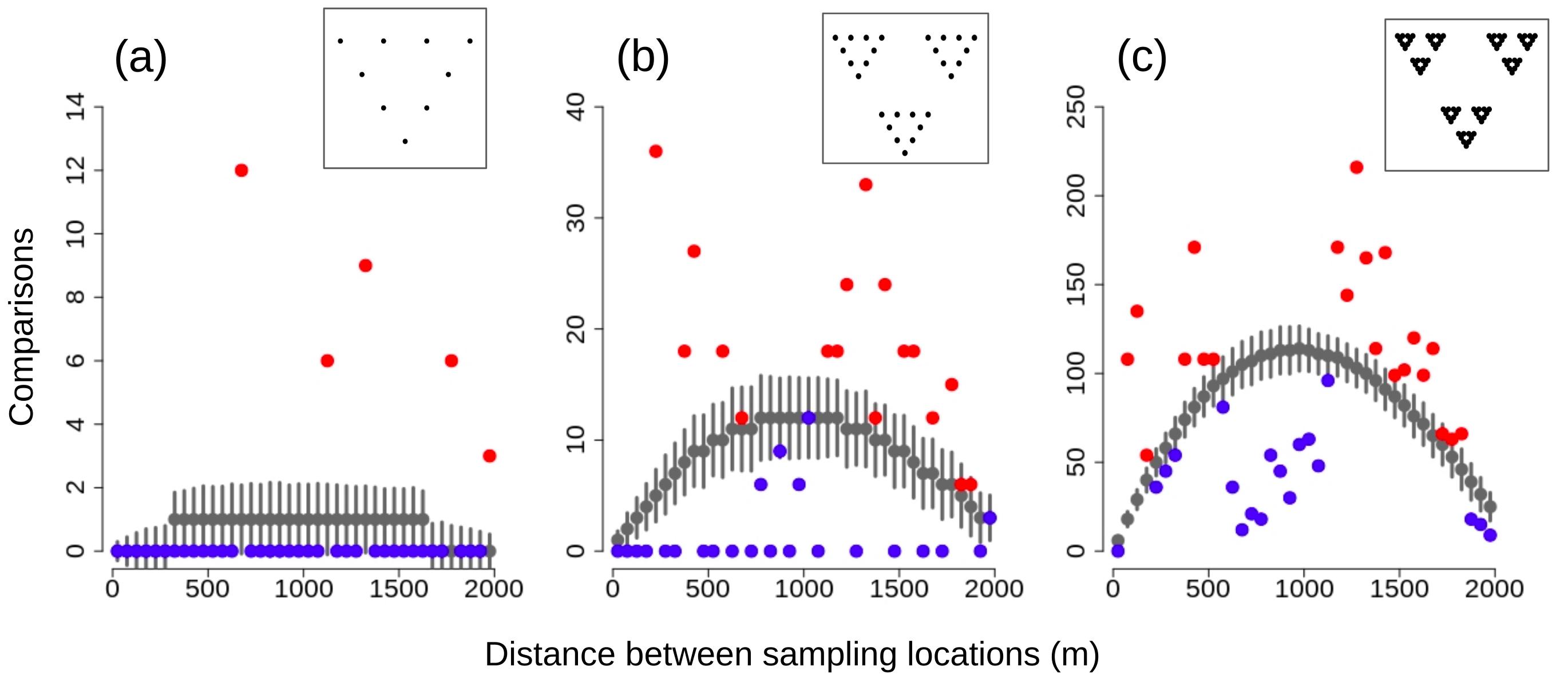

### fbsd-fig3-2017-supplement.jpeg

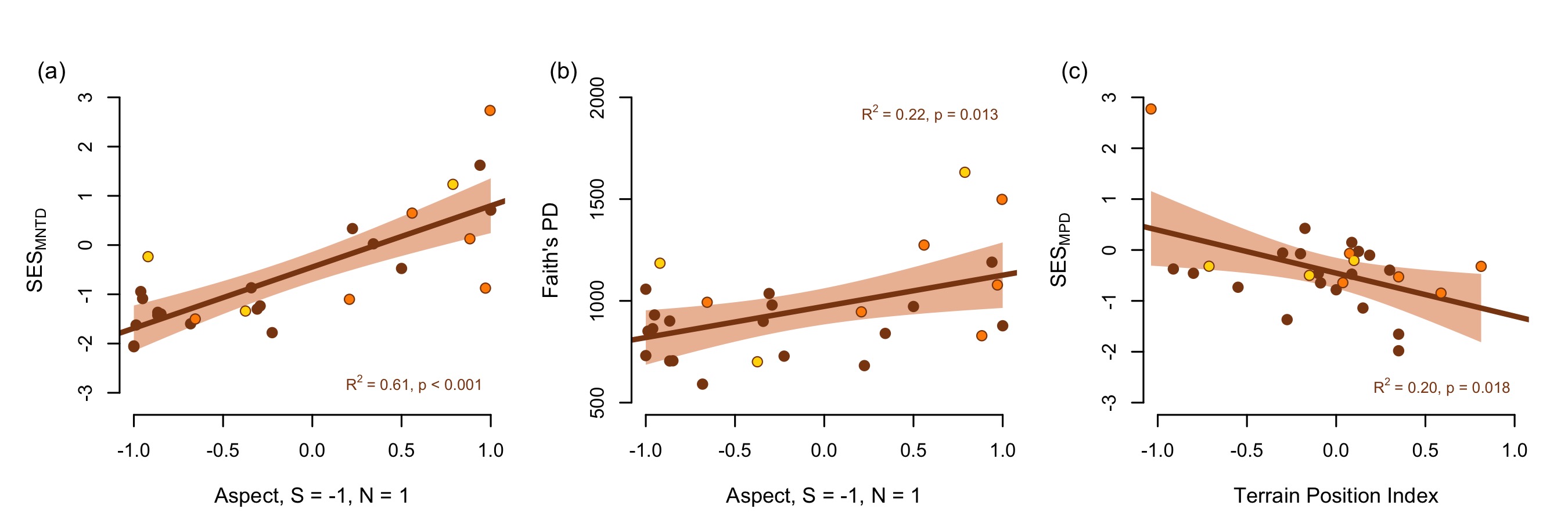

### fbsd-fig3-supplement.jpeg

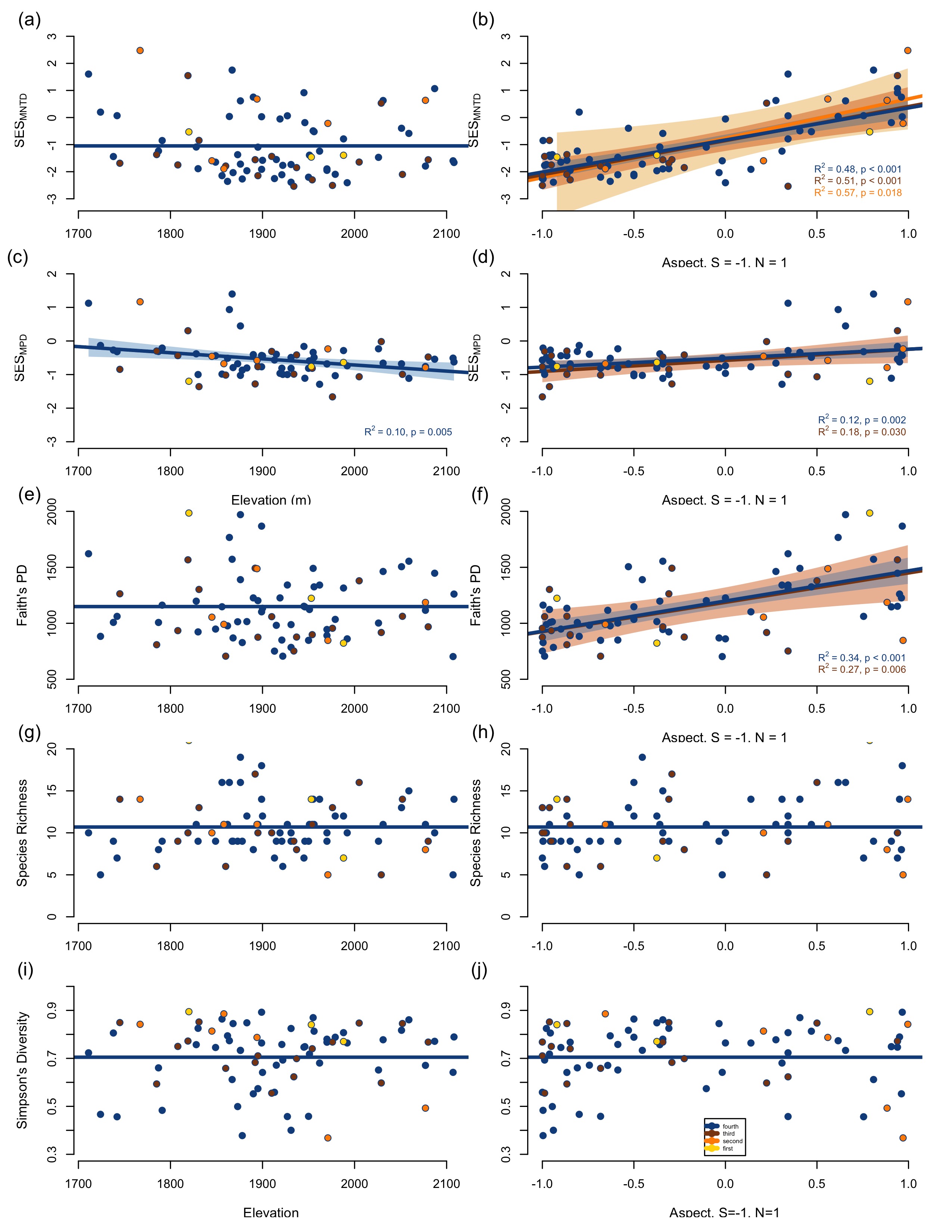

### fbsd-fig3.jpeg

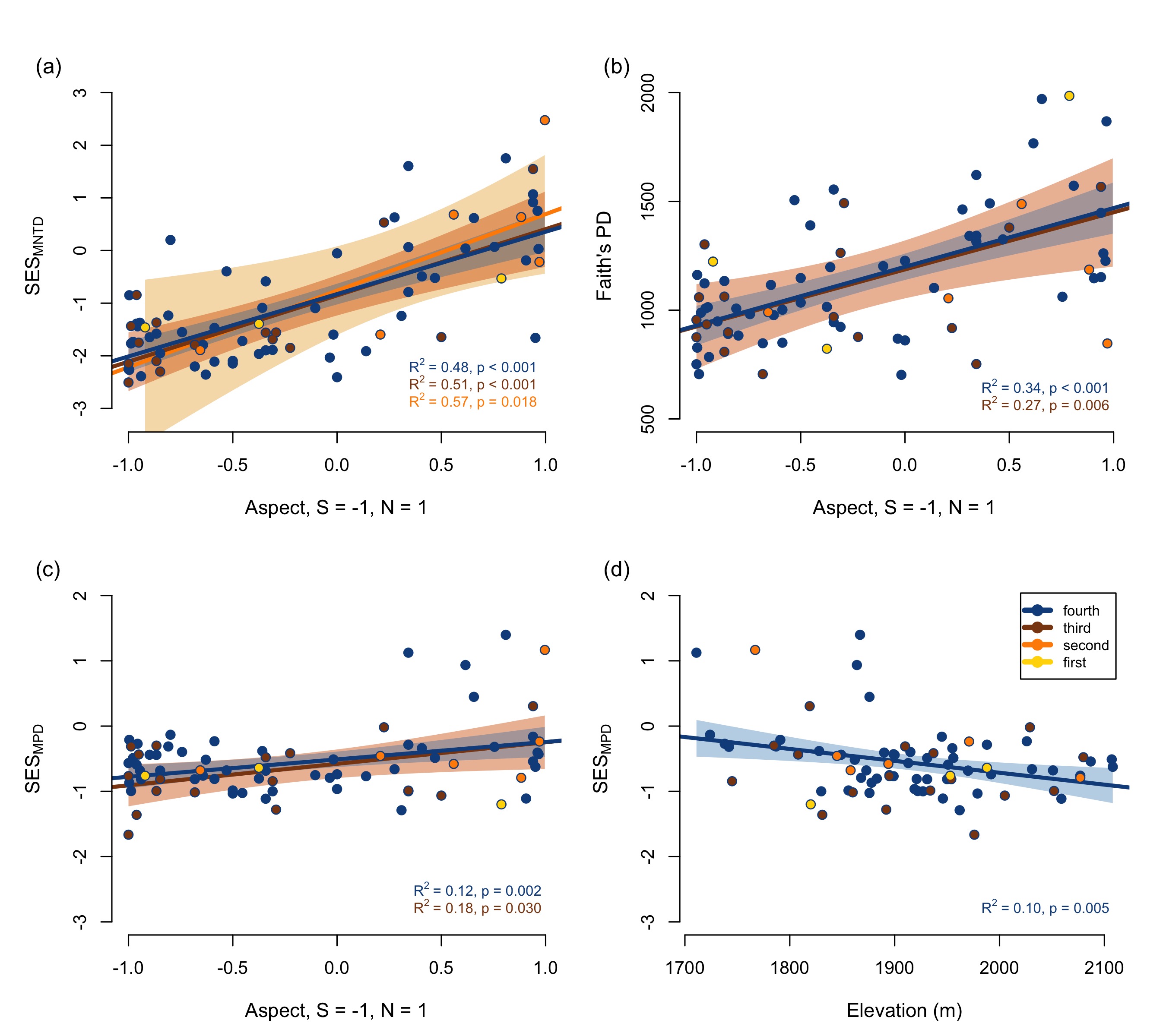

### fbsd-fig4-2017-supplement-raw.jpeg

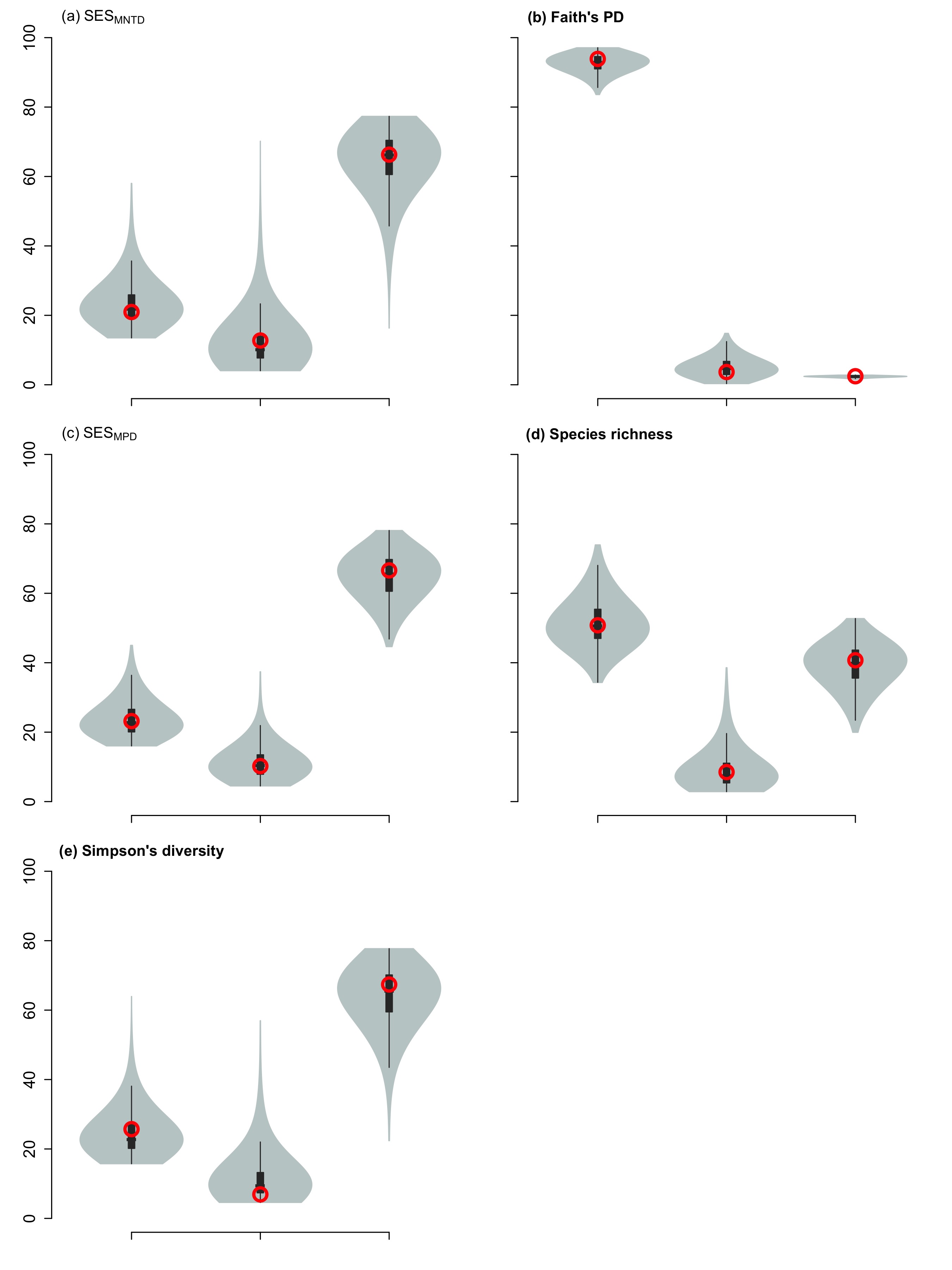

### fbsd-fig4-2017-supplement.jpeg

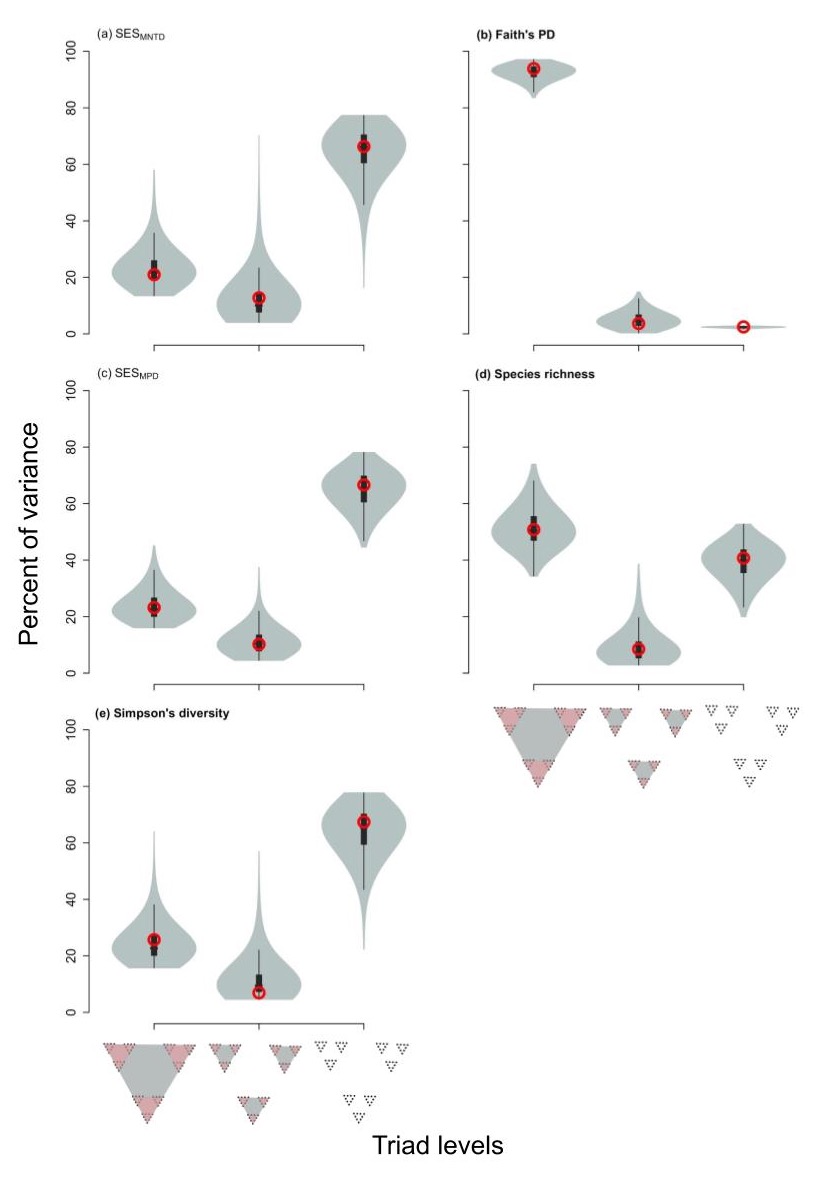

### fbsd-fig4-raw.jpeg

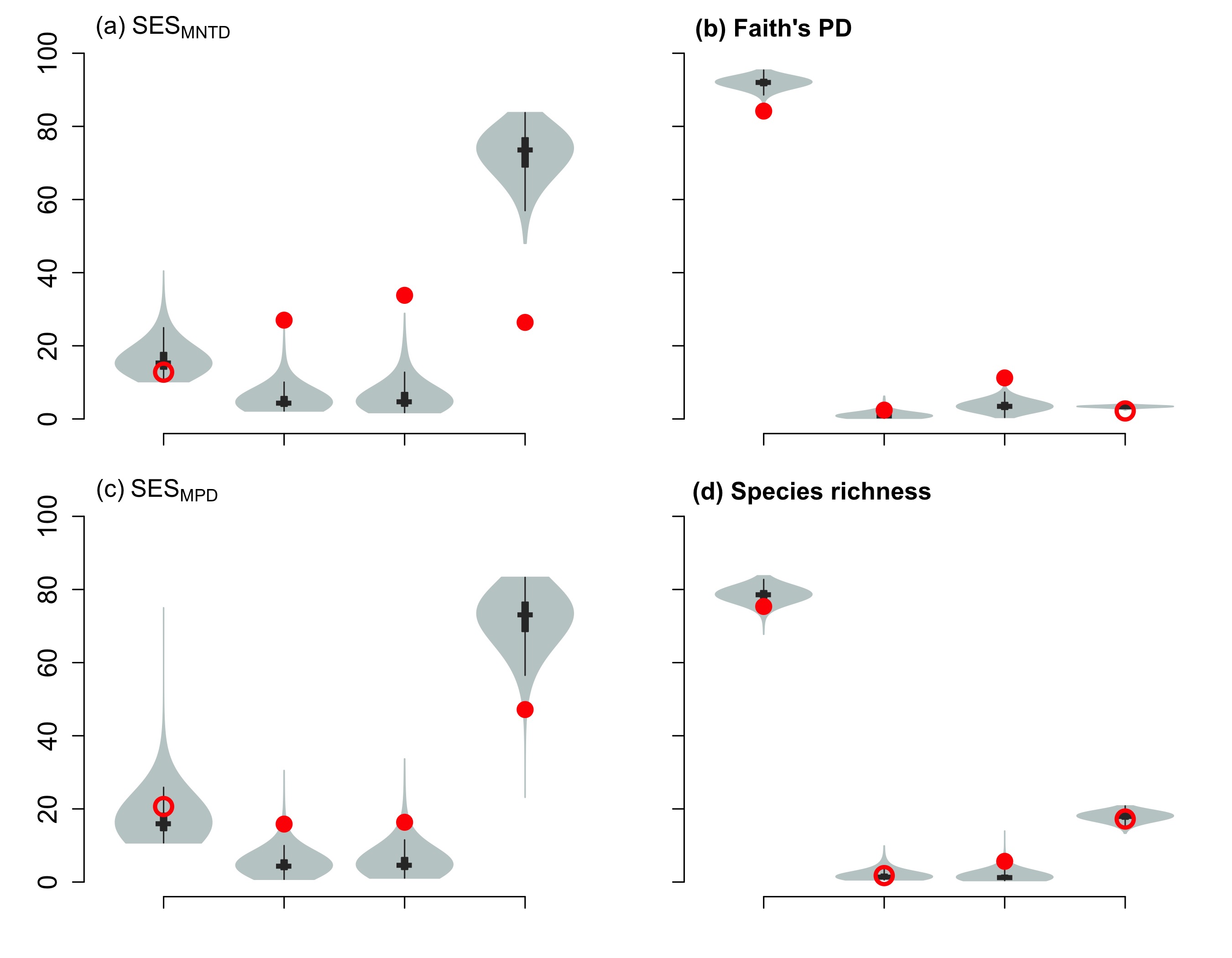

### fbsd-fig4-supplement-raw.jpeg

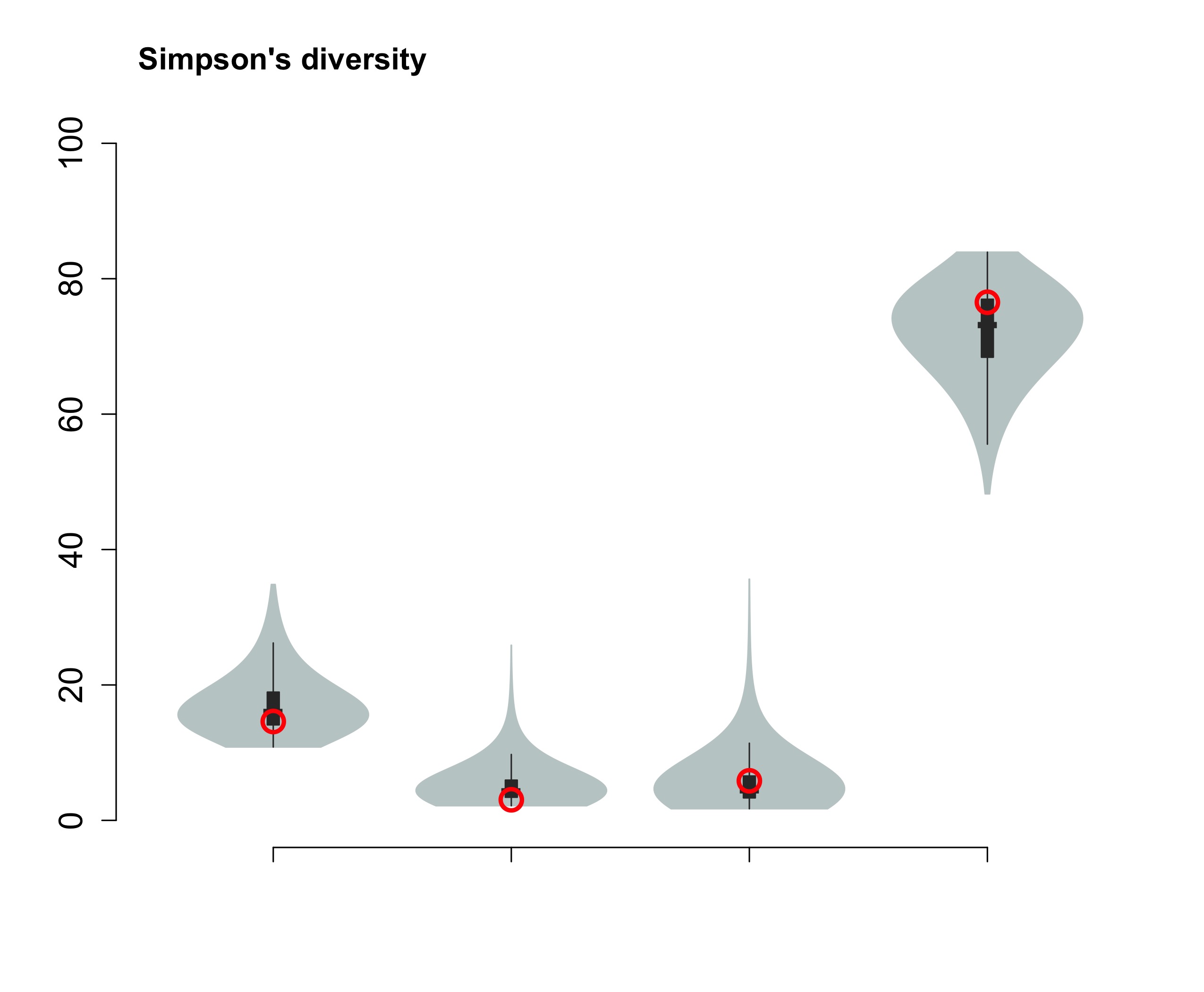

### fbsd-fig4-supplement.jpeg

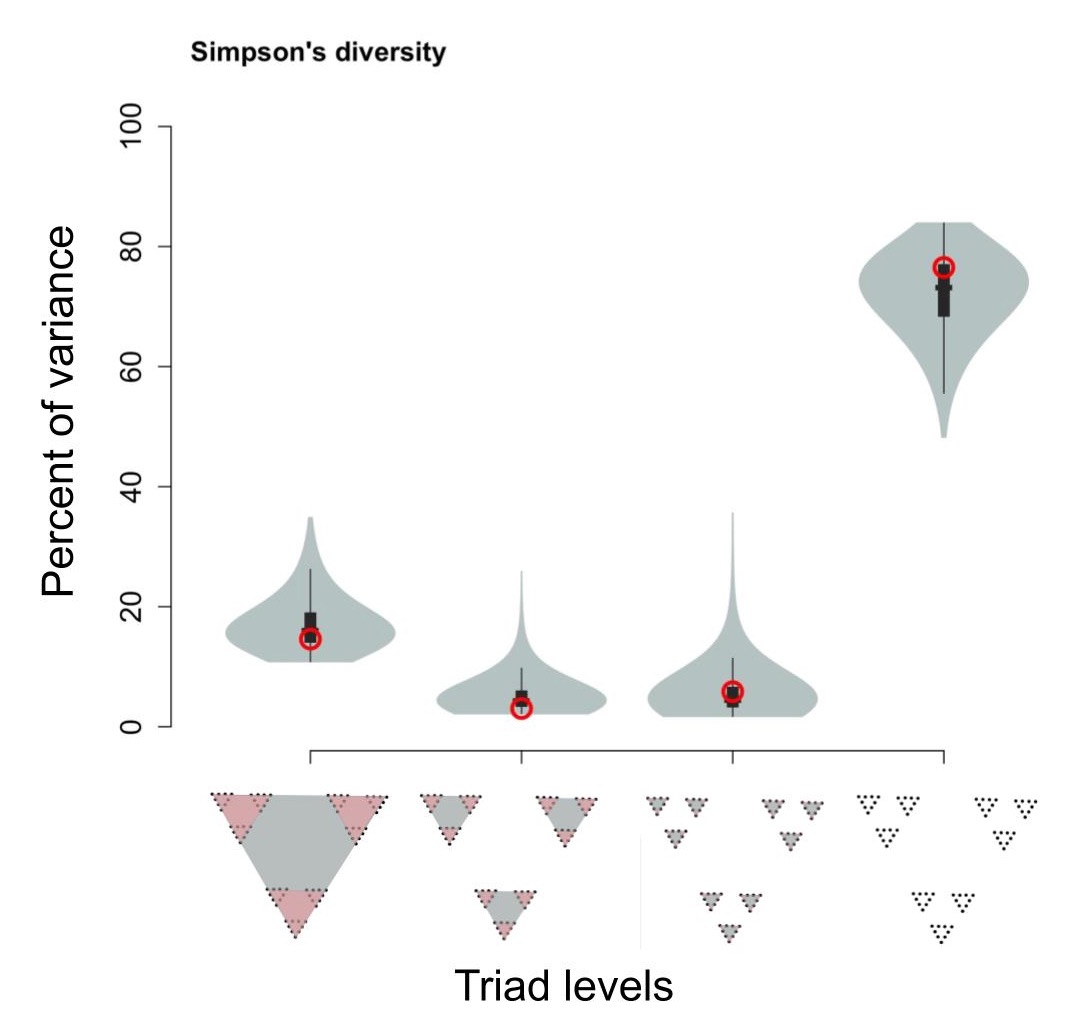

### fbsd-fig4.jpeg

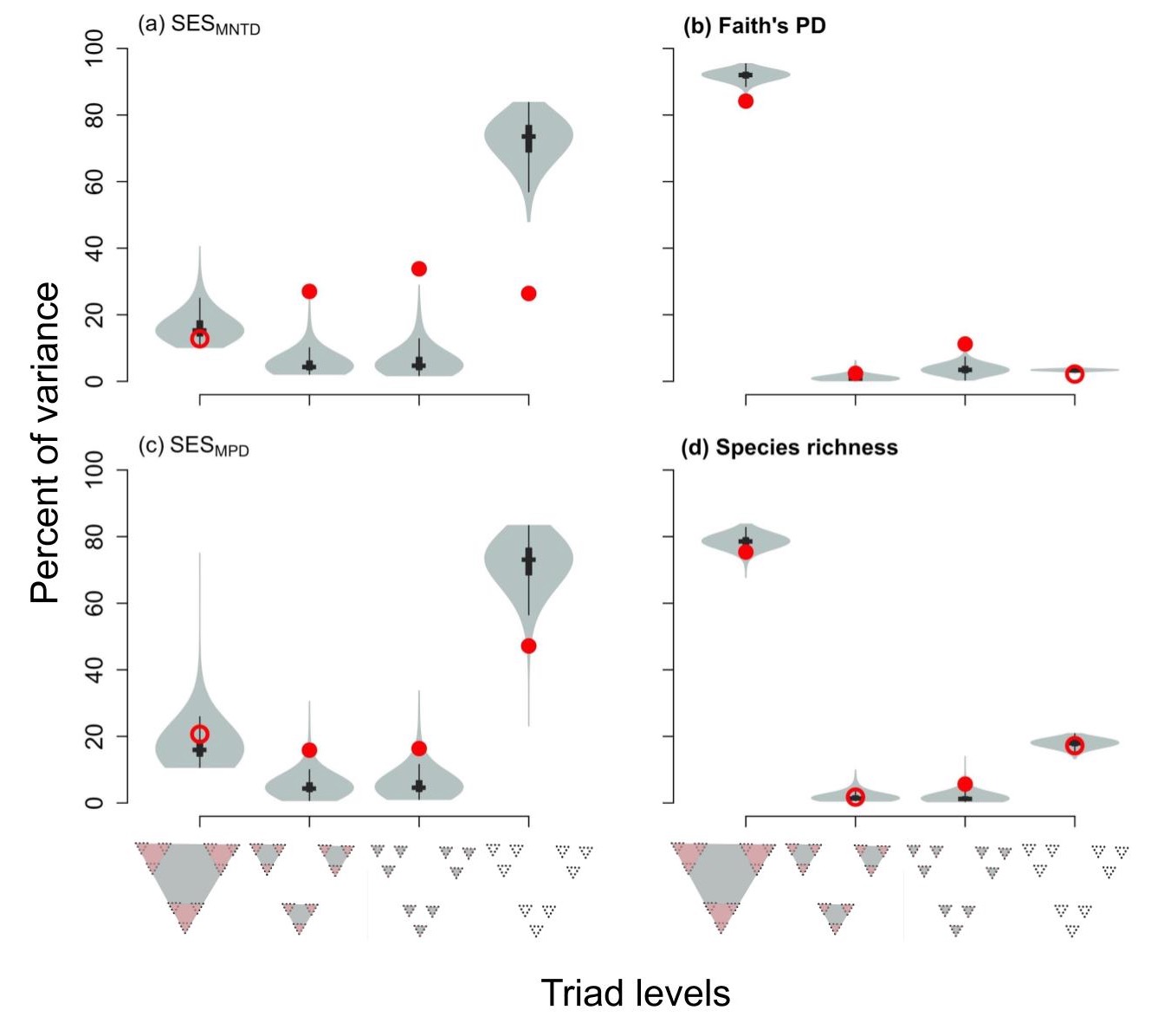
